# Cathepsin g degrades synovial fluid lubricin: relevance for osteoarthritis pathogenesis

**DOI:** 10.1101/792184

**Authors:** Shan Huang, Kristina A. Thomsson, Chunsheng Jin, Sally Alweddi, André Struglics, Ola Rolfson, Lena I Björkman, Sebastian Kalamajski, Tannin A. Schmidt, Gregory D. Jay, Roman Krawetz, Niclas G. Karlsson, Thomas Eisler

**Author notes:** Corresponding author - Niclas G. Karlsson, Department of Medical Biochemistry and Cell Biology, Institute of Biomedicine, Sahlgrenska Academy, University of Gothenburg, Sweden. Medicinaregatan 9A, 405 30 Gothenburg, Sweden. These authors have contributed equally to this work (Shan Huang, Kristina A. Thomsson). These authors have jointly supervised this work (Niclas G. Karlsson, Thomas Eisler).

## Abstract

Lubricin (PRG4) is a mucin type protein that plays an important role in maintaining normal joint function by providing lubrication and chondroprotection. Improper lubricin modification and degradation has been observed in idiopathic osteoarthritis (OA), while the detailed mechanism still remains unknown. We hypothesized that the protease cathepsin G (CG) may participate in degrading lubricin in synovial fluid (SF). The presence of endogenous CG in SF was confirmed in 16 patients with knee OA. Recombinant human lubricin (rhPRG4) and native lubricin purified from the SF of patients were incubated with exogenous CG and lubricin degradation was monitored using western blot, staining by Coomassie or Periodic Acid-Schiff in gels, and with proteomics. Full length lubricin (∼300 kDa), was efficiently digested with CG generating a 25-kDa protein fragment, originating from the densely glycosylated mucin domain (∼250 kDa). The 25-kDa fragment was present in the SF from OA patients, and the amount was increased after incubation with CG. A CG digest of rhPRG4 revealed 135 peptides and 72 glycopeptides, and confirmed that the protease could cleave in different domains of lubricin. Our results suggest that synovial CG may take part in the degradation of lubricin, which could affect the lubrication of OA joints.

## Introduction

Osteoarthritis (OA) is the most common degenerative joint disease, which affects people worldwide^1^. OA involves a progressive destruction of cartilage, bone and ligaments, sometimes causing extensive pain, reduced joint flexibility and may lead to severely impaired quality of life and rising health care costs. Recent research emphasises idiopathic OA as a multifactorial joint disease^2^ with an increasing number of studies demonstrating a low-grade inflammation with an accumulation of pro-inflammatory cytokines in synovial fluid (SF), which is associated with the pathological process of OA^3^. Cytokine triggered enzymatic degradation of matrix proteins is believed to contribute to the cartilage erosion found in OA^4-7^.

Lubricin, also known as Proteoglycan 4 (PRG4) is a large (∼ 300 kDa) extensively *O*-linked glycosylated mucinous protein in SF, and plays a critical role in maintaining cartilage integrity by providing boundary lubrication and reducing friction at the cartilage surface^8^. Besides the principle function of lubrication, lubricin also has growth-regulating properties^9^, prevents cell adhesion, provides chondroprotection^10-12^, and plays a role in the maturation of the subchondral bone^13^. Lubricin is predominantly synthesised and expressed by superficial zone chondrocytes at the cartilage surface layer^14^, but can also be secreted into the SF by synovial fibroblasts^15^ and stromal cells from peri-articular adipose tissues^16^. More recently it was reported that other sites and tissues, such as tendons, liver, kidney, skeletal muscles, and the ocular surface, also express lubricin^17-19^.

In the joint, lubricin is an extended molecule existing both as monomers and ashigher molecular mass complexes^18^. Lubricin has an approximately equal mass proportion of protein and oligosaccharides, and contains one central mucin-like region, two somatomedin B homology domains (SMB) in the *N*-terminal, one heparin or chondroitin sulfate binding domain, and one hemopexin-like domain (PEX)^17,20^ in the *C*-terminal (Fig. 1). The multiple domain structure contributes to diverse biological roles of lubricin. The *O*-linked oligosaccharides in the mucin domain render lubricin a low friction brush-like construct with repulsive hydration forces, provides lubrication during boundary movement, and makes lubricin water-soluble in SF^21^. The oligosaccharides are linked to abundant Thr/Ser residues, which are found in the repeat region, consisting of repeats with minor variations of the amino acid sequence ‘EPAPTTPK’, as well as its flanking, non-repeating Thr/Ser rich regions. The sparingly glycosylated terminal regions SMB and PEX domains have in other proteins been demonstrated to regulate innate immune processes by interacting with both complement and coagulation factors ^22^ and assisting matrix protein binding^23^. For over a decade, researchers have demonstrated that lubricin plays a vital role in inflammatory joint disease. Prevention of lubricin degradation may serve as a therapy in an early stage inflammatory arthritis^24-28^. Altered lubricin structure and change of lubricin concentrations which disrupts cartilage boundary lubrication is found in SF of OA patients^29^. As such, lubricin degradation may play a role in OA associated inflammation. A limited number of proteases have been identified to degrade lubricin, among the more studied are lysosomal cysteine proteases cathepsin B, S, L and neutrophil elastase^26,30,31^. Cathepsin G (CG) is one of the major neutrophil serine proteases that is synthesised in bone marrow and subsequently stored in the azurophil granules of polymorphonuclear neutrophils^32,33^. Upon activation of the granulocytes, CG is released at sites of inflammation and plays a crucial part in degrading chemokines and extracellular matrix (ECM) proteins, as well as regulating and activating pro-inflammatory cytokines^34^. Therefore, CG is reported to be active in various chronic inflammatory diseases^35^ and has for instance been shown to be highly active in the SF of rheumatoid arthritis patients.

**Figure 1.**
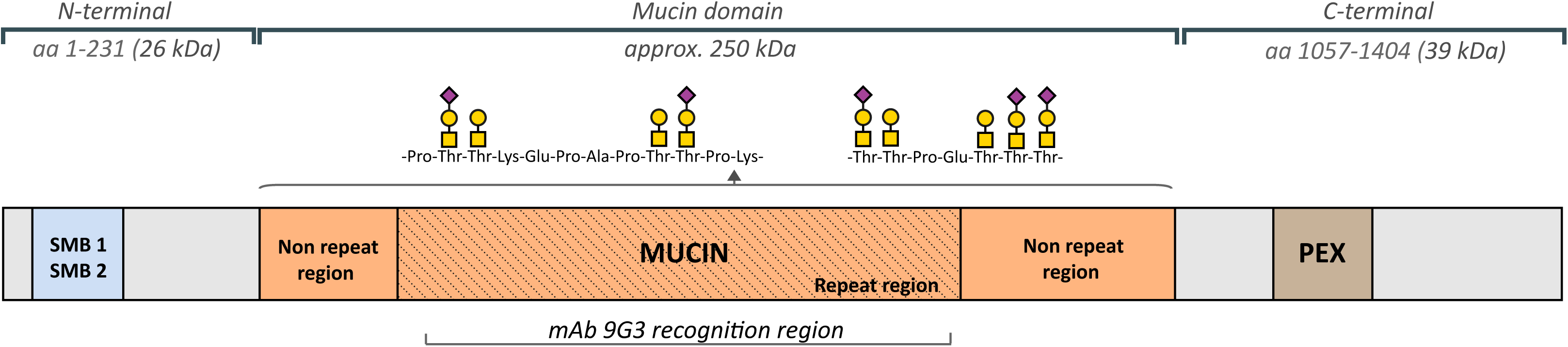
**(a)** Description of lubricin and its domains. Lubricin contains a heavily glycosylated Pro/Thr rich mucin domain, flanked by two somatomedin B domains (SMB) and a hemopexin (PEX) domain. The *O*-glycans are linked to Thr and Ser residues, and increases the size of the mucin domain from 86 kD (without glycans) to approximately 250 kD. The monoclonal lubricin antibody 9G3 has been shown to bind the glycosylated peptide sequence ‘KEPAPTTT’ which is found eight times in the repeat region. Monosaccharides constituting lubricin oligosaccharides are represented by Symbol Nomenclature for Glycans (SNFG)^53^, where *N*-acetylgalactosamine (GalNAc) is a yellow square, galactose (Gal) is a yellow circle and *N*-acetylneuraminic acid (NeuAc) is a purple diamond.

Despite evidence indicating that lubricin modification contributes to some extent to OA initiation and pathological development, research on and understanding of enzymatic degradation of lubricin in OA is still limited. Hence, in this project, the serine protease CG was investigated for its participation in lubricin degradation in relation to OA. The presence of endogenous CG was detected in SF samples from 16 patients with idiopathic knee OA. The potent lubricin degrading ability of CG was confirmed both *in vitro* and *in vivo*, and the CG generated fragments of recombinant lubricin were identified.

## Results

### Identification of CG in synovial fluid of OA subjects

CG has previously been shown to be present in OA SF, however in a low abundance ^36^. Since we hypothesized that CG plays a role in the OA pathology, we verified in our SF sample collection that CG was present. SF samples were collected from 16 late-stage OA patients and analysed for the presence of CG using SDS-PAGE and western blot with a polyclonal anti-CG antibody. We could detect CG in all the SF samples at varying concentrations, the results are presented relative to a reference sample which was analysed on the same gel (Fig. 2a,b and Supplementary Fig. S1). Our results revealed that the CG levels in OA SF were estimated to be in the low ng range (1-10 ng) per μl SF. This is in line with a previous study using a general colorimetric substrate assay for trypsin/chymotryptic-like proteases in OA and RA SF^36^.

**Figure 2.**
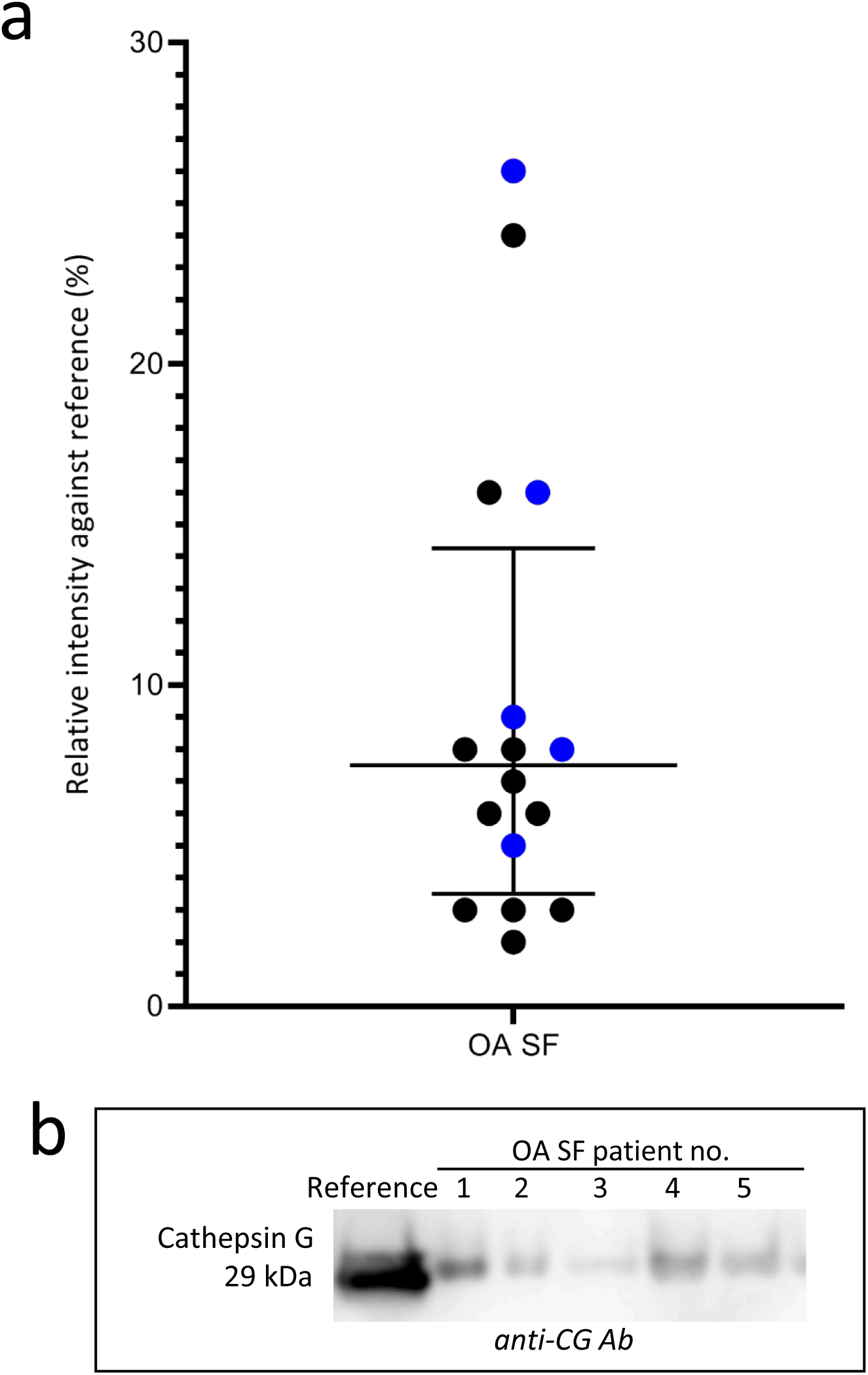
Endogenous Cathepsin G (CG) in synovial fluid (SF) from OA patients. **(a)** CG was detected in SF from 16 OA patients. SF was analysed with SDS-PAGE followed by western blot using a polyclonal anti-CG antibody. Bands corresponding to CG were detected at 29 kDa, and the results are displayed as relative intensities (% of absorbance), compared to a reference compound analysed on the same gel (22 ng CG). Individual subject values from a selected region from the western blot showing the 29 kDa CG bands from five patients **(b)** are highlighted in blue. The full-length western blot is displayed in Supplementary Fig. S1.

### CG degrades recombinant and native lubricin

In order to investigate if CG was able to digest lubricin, we performed incubations using recombinant lubricin (rhPRG4), the degradation products were separated with SDS-PAGE and detected with lubricin mAb 9G3. This antibody targets the glycosylated mucin domain^19^. The results indicated that rhPRG4 was indeed degraded after extended incubation with only one major degradation product detected at approximately 25 kDa (Fig. 3a).

**Figure 3.**
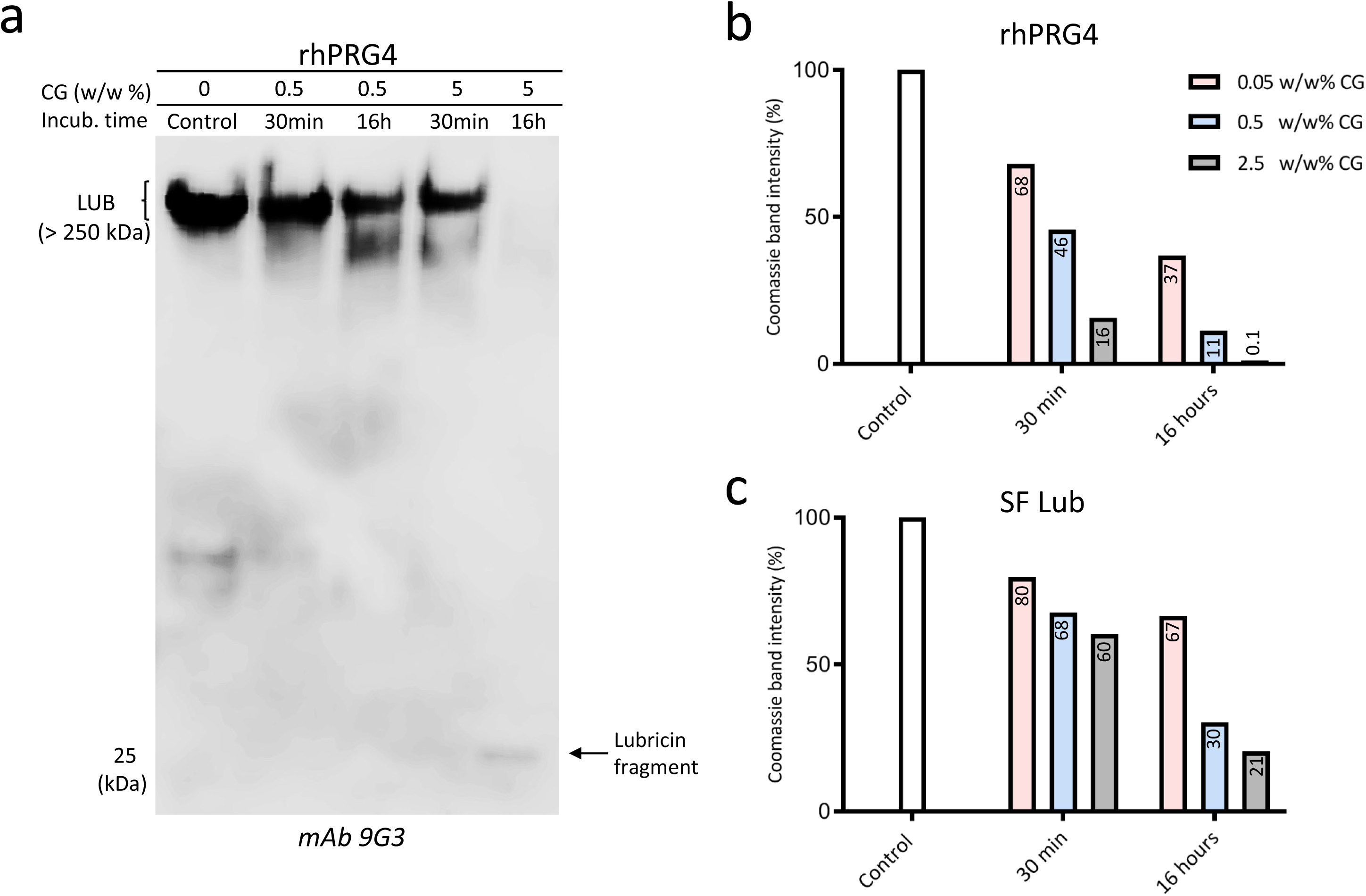
Cathepsin G (CG) degrades lubricin. rhPRG4 and purified lubricin from synovial fluid (SF Lub) were incubated with CG or without CG (control) at increasing enzyme to substrate ratios (weight/weight) in PBS for 30 min or 16 hours, followed by analyses on tris-acetate gels (3-8%). rhPRG4 was detected with western blot using mAb 9G3 **(a)** or using Coomassie Blue stain (rhPRG4 and purified SF lubricin, Supplementary Figs S2 and S3). Coomassie Blue band intensities of full length lubricin incubated with CG compared to the control lubricin band (no CG) are displayed in **(b)** and **(c)** with the relative intensities in % displayed as a number in each bar.

To investigate if there was a difference in the ability to digest two different glycoforms of lubricin, we monitored the effect of CG digestion of both rhPRG4 and native lubricin using a Coomassie protein staining.Both rhPRG4 and native lubricin purified from SF were found to be degraded in a dose-dependent manner. No major degradation products were observed, and the degradation increased with prolonged incubation time (Figs 3b, c and Supplementary Figs S2, S3). rhPRG4 was found to be efficiently degraded by CG already after 30 minutes of incubation, illustrated by that 32%, 54% and 84% of the full length protein had been degraded, with increasing CG-to-protein ratio. After 16 hours, rhPRG4 was completely degraded (Fig. 3b). Native lubricin was more resilient to CG digestion. After 16 hours of incubation with the higher CG concentrations (0.5 and 2.5 w/w % CG), 30 and 21 % of the full length lubricin remained undigested, respectively. (Fig. 3c). These results suggested that the protein core of native lubricin was less accessible for CG digestion. We also showed that the effect of CG degradation of purified SF lubricin could be totally abolished using CG inhibitors (Supplementary Fig. S4). In all, we concluded that CG was capable of degrading lubricin both in its recombinant and native form.

### CG degrades lubricin mucin domains in the presence of SF

The decrease in Coomassie stain of lubricin after incubation with CG (Figs 3b, c and Supplementary Figs S2, S3) indicated that the protein backbone was substantially effected. We were interested to see if CG also could hydrolyse peptide bonds within the highly glycosylated mucin domains. Mucin domains are renowned for being hard to digest with proteolytic enzymes. Recombinant (rhPRG4) and purified lubricin from SF were incubated for two hours with CG, followed by separation on gels and developed with Periodic Schiff’s base stain. This stains the glycans on the heavily glycosylated mucin domain of lubricin. Only two glycosylated degradation products (15- and 25-kDa) were detected on the SDS-gels (Fig. 4a), indicating that additional proteolytic products were of lower mass. The result supported previous observations, where both the glycosylated and non-glycosylated regions of lubricin were proteolytically digested by CG.

**Figure 4.**
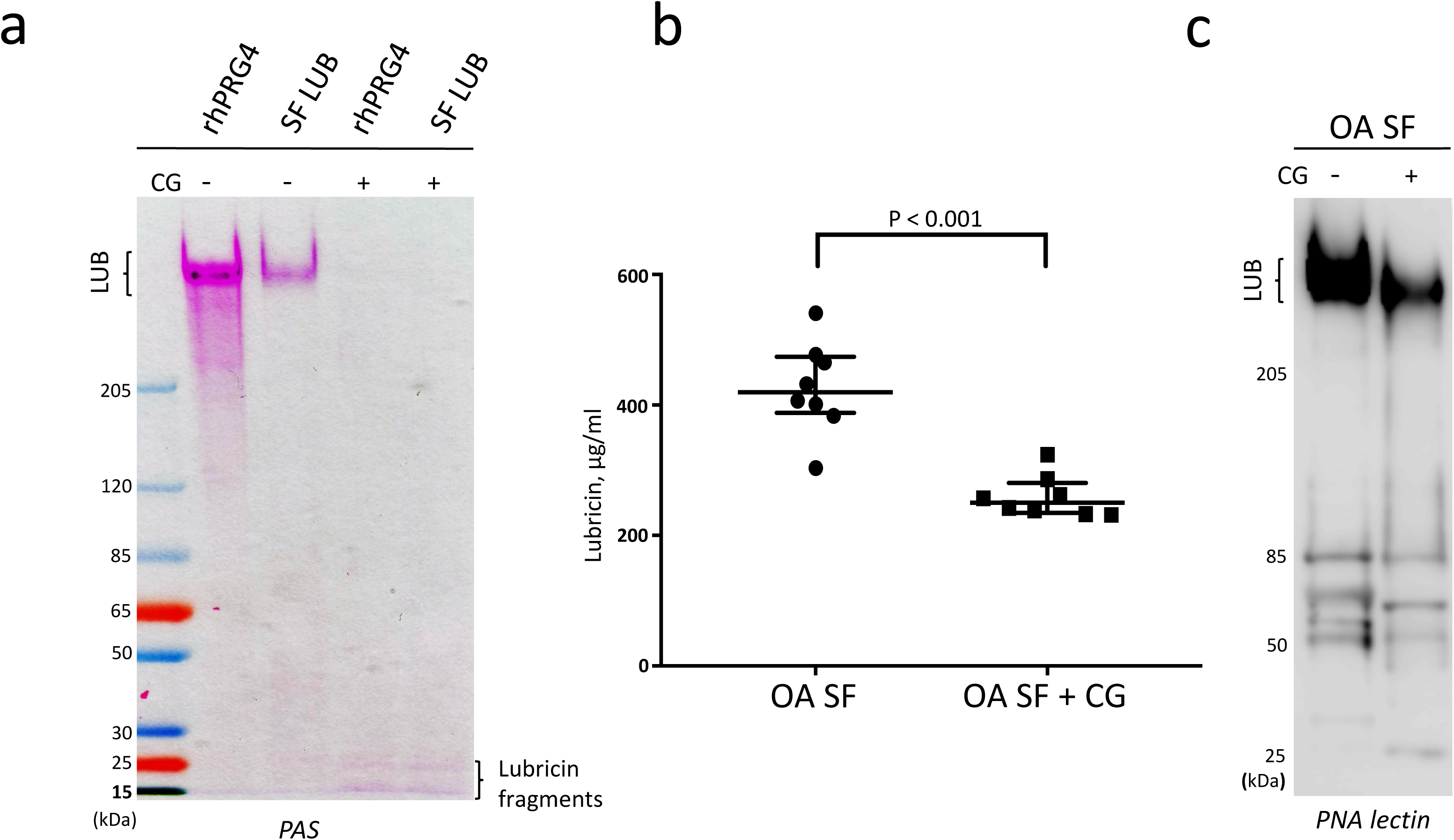
PAS and PNA lectin detection of lubricin and glycosylated lubricin fragments after incubation with Cathepsin G (CG). **(a)** Periodic Acid Schiff (PAS) staining of recombinant lubricin (rhPRG4) and purified lubricin from synovial fluid (SF LUB). rhPRG4 and SF LUB were incubated with 5 w/w% CG (enzyme to substrate ratio by weight) in PBS for 2 hours, followed by analysis with SDS-PAGE and staining with PAS, which stains densely glycosylated proteins. **(b)** Lubricin quantification in SF using ELISA. SF (2 μl) from eight OA patients were incubated with CG (44ng) or without CG for 2 hours at 37°C. Lubricin was measured using a sandwich ELISA, with mAb 9G3 as the catching Ab and *Peanut agglutinin* (PNA) lectin for detection, and compared to a standard curve with rhPRG4. Data are presented as mean +/- SEM. **(c)** PNA lectin western blot of SF incubated with CG. SF (2.5 μL) from an OA patient was incubated with or without CG (55 ng) for 2 hours at 37°C.

The influence of other components present in SF on CG degradation of the lubricin mucin domain was also assessed. Eight OA SF samples were incubated with or without supplement of exogenous CG. Lubricin was quantified using ELISA with mAb 9G3 as the catching antibody and PNA lectin for detection. Peanut agglutinin lectin (PNA) binds to terminal galactose residues that are abundantly expressed on lubricin^17^. With this ELISA method, we could estimate the amounts of lubricin in OA SF to be in the range of 300-550 µg/ml (Fig. 4b). After incubation with CG (8-15 w/w%, enzyme to lubricin weight ratio), the amount of lubricin was decreased approximately 50%. This experiment highlights that lubricin can be degraded in SF by an exogenous supply of CG. CG degradation of lubricin was less efficient in the presence of SF, compared to when performed in PBS (Fig 4a and Supplementary Fig. S5).

CG degradation of the lubricin mucin domain in SF was also visualized using the PNA lectin by western blots. After two hours of incubation of OA SF with CG, the intensity of the band corresponding to full length lubricin had decreased considerably compared to the control without CG, and the formation of a weakly stained band at 25 kDa was observed (Fig. 4c). Western blot of SF incubated without CG revealed that in addition to full length lubricin, several other PNA reactive components of unknown origin at 50-100 kDa was detected. These components probably represent other PNA reactive proteins in SF as well as degradation products of lubricin.

### The 25-kDa lubricin fragment is found in SF from OA patients and is increased after CG incubation

We were interested to investigate if the 25 kDa mucin fragment generated from lubricin also could be found in SF, providing evidence that lubricin is degraded in its natural environment. Indeed, by increasing exposure time of the western blot using the mucin domain mAb 9G3, a single lubricin fragment at 25 kDa of endogenous origin was observed in SF from 13 OA patients without CG supplementation (Figs 5a,b and Supplementary Fig. S6). The selected region from a western blot shows the lubricin degradation fragment at 25 kDa in SF from six patients (Fig. 5a).

**Figure 5.**
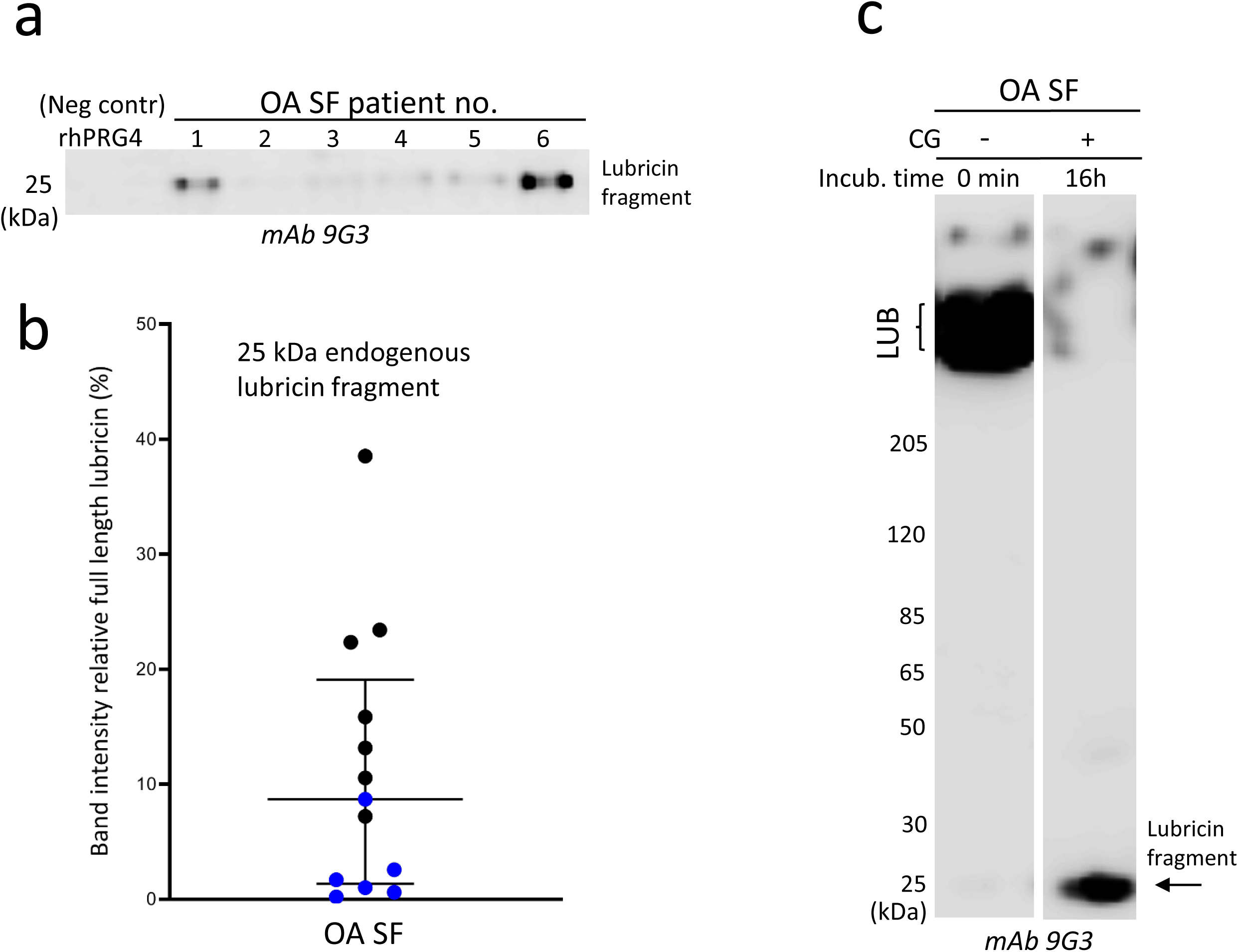
Identification of the 25 kDa glycosylated lubricin fragment from synovial fluid (SF) of OA patients. **(a)** Detection of an endogenous lubricin degradation fragment at 25 kDa in SF from OA patients. SF samples (2 μl) were analysed with SDS-PAGE, followed by western blot using mAb 9G3. Selected region from a western blot showing examples of a lubricin degradation fragment at 25 kDa from six OA patients. The full-length western blot is displayed in Supplementary Fig. S6. **(b)** Semiquantification of the 25-kDa lubricin fragment in SF from 13 OA patients. The 25-kDa band intensities from western blot analyses are plotted relative to the full length lubricin band for every patient sample. rhPRG4 (1 μg) was used as negative control. Individual subject values from the western blot in (a) are highlighted in blue. **(c)** SF incubated with exogenous CG. SF (2 μL) from an OA patient was incubated with CG (44 ng) for 0 or 16 hours, followed by SDS-PAGE and detection with mAb 9G3. The original western blot is displayed in Supplementary Fig. S7.

We could also show that the amount of the 25 kDa fragment was increased after addition of exogenous CG to SF (Fig. 5c and Supplementary Fig. S7). Complete degradation of full length lubricin was accompanied by a concomitant increase of a 25-kDa lubricin fragment, similar as was observed for rhPRG4 (Fig. 3a). We found that extending the incubation of lubricin with CG even for a longer period (> 16 hours, results not shown) eventually decreased the intensity of the 25-kDa fragment, indicating that CG was capable of digesting this component even further.

These experiments suggest that endogenous SF CG is at least partly responsible for generating the 25-kDa lubricin fragment found in OA patients.

### Proteomics and glycoproteomics analyses of rhPRG4 digested with CG

In order to identify CG cleavage sites within lubricin, rhPRG4 was incubated with CG in PBS overnight at 37°C at a ratio of 1:45 (enzyme to substrate, by weight), and the obtained degradation products analysed with LC-MS/MS. We identified 135 non-glycosylated peptides and 72 glycopeptides in the size range of 6-37 amino acids, the majority from the *N*- and *C*-terminal of lubricin (Supplementary Tables S1 and S2). CG has been described to have a combination of tryptic and chymotryptic type specificity, but is also reported to cleave at other amino acids^37-39^. An overview of the cleavage sites in lubricin detected here are displayed in Fig. 6a. As a control experiment, we performed semi-tryptic searches of tryptic digests of rhPRG4, which revealed that 13 peptides could originate from other sources than CG digestion, for example autoproteolytic degradation. The most frequent cleavage site was C-terminal of lysine residues (33 %), a site which has been reported previously for CG^37^. Many peptides were found to be overlapping within the same regions in the peptide backbone. CG cleavage sites within the glycosylated mucin domain were identified after manual evaluation of MS spectra.

**Figure 6.**
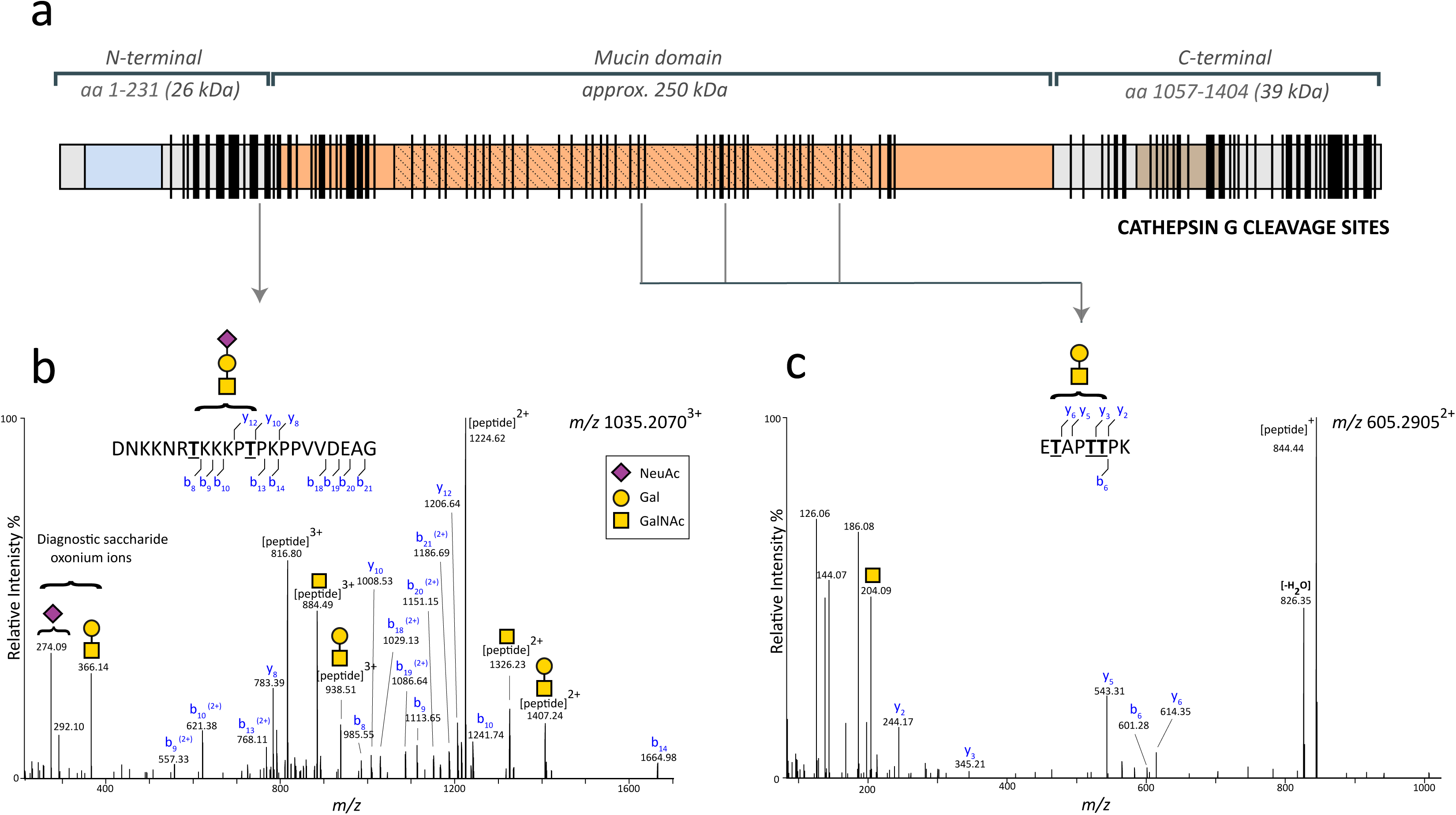
**(a)** Recombinant lubricin (rhPRG4) was digested with cathepsin G (CG), followed by analyses of the peptides with proteomics and glycoproteomics. Identified cleavage sites along the lubricin protein are represented by vertical lines. The complete peptide/glycopeptide lists are found in Supplementary Tables S1 and S2. **(b)** and **(c)** Higher-energy collisional dissociation (HCD) spectra of *O*-glycopeptides of rhPRG4 digested with CG, and analysed with LC/MS/MS. Diagnostic glycan oxonium ions are detected in the lower mass range (*m/z* 100-400). b/y-ions are detected without glycan substituents. Potential *O*-glycan sites (Ser/Thr) are underlined and in bold. **(b)** The glycopeptide at *m/z* 1035.2070 (3+) eluting at 14.3 min was deduced to have the sequence ‘DNKKNRTKKKPTPKPPVVDEAG’ and carry the Sialyl T-antigen (NeuAc-Gal-GalNAc). The glycopeptide originates from the lubricin N-terminal (aa positions 202-233). **(c)** The glycopeptide at *m/z* 605.2905(2+) eluting at 15.0-15.4 min was deduced to have the sequence ‘ETAPTTPK’ and carry the T-antigen (Gal-GalNAc). The peptide originates from the mucin domain and is found three times in lubricin (aa positions 615-622,703-710, 825-832).

The glycopeptides constituted glycoforms made up of 35 peptide cores, carrying glycans of different monosaccharide compositions, and consisting of Hex, HexNAc and NeuAc residues, matching simple core 1 type *O*-glycosylation that is produced in CHO cells from where the rhPRG4 was expressed (Supplementary Table S2). Higher-energy collisional dissociation (HCD) fragmentation did not reveal glycan site specific information, however all but one deduced peptide contained potential *O*-glycan sites (Thr or Ser, and less common, Tyr). The peptide D.MDYLPRVPN.Q from the C-terminal (aa 1122-1130) was detected as two glycoforms and proposed to be glycosylated on a Tyr residue (Supplementary Fig. S8). The peptide L.RNGTVLAF.R (aa 1158-1165) was detected as two glycoforms, revealed monosaccharide compositions matching that of *N*-glycans of high mannose type (Supplementary Fig. S9). We have also detected this potential *N*-glycan site in tryptic digests (data not shown). Mass spectra of three glycopeptides with the peptide sequences ‘ETAPTTPK’, ‘KEPAPTTPK’ and ‘KEPAPTTPKKPAPK’ from within the mucin repeat region are shown in Fig 6c and Supplementary Figs S10 and S11, respectively. These repeat peptides are found 3, 17, and 2 times respectively in the mucin domain. Together with the identification of two nonglycosylated repeat peptides (‘KEPAPTTPKKPAPK’ and ‘SAPTTTKEPAPTTTK’, Supplementary Table S1), these data provided mass spectrometric evidence for that CG may cleave within the mucin region. The N-terminal derived glycopeptide with the sequence ‘K.DNKKNRTKKKPTPKPPVVDEAG.S’ (aa 202-223) contains two potential glycosylation sites (Thr208 and Thr213) and was detected as two glycoforms, where the monosaccharide compositions of the first glycoform supports the presence of a T-antigen (Galβ1-3GalNAc), the second the sialylated T-antigen (Fig. 6b). CG was also able to cleave this glycopeptide stretch further (Supplementary Table S2). The CG cleavage sites within amino acid sequence 202-223 are summarized by vertical lines:

DN|K|K|NRTK|KKPTPKPPVV|D|E|A|G. This example illustrates that CG can cleave after many different amino acids in lubricin, however at low efficiency. This promiscuity in cleavage site specificity has been highlighted in studies with defined substrates^37^.

## Discussion

Despite being one of the most common and disabling joint diseases, OA pathogenesis and development are still largely unknown. The success of introducing recombinant lubricin for OA treatment in animal models^40,41^, indicate a role of lubricin in OA pathogenesis. To date, improper lubricin post-translational modifications, including glycosylation profile change^42^, forming complex with other matrix proteins^43^, and proteolytic cleavages are all reported in OA. This may ultimately alter the surface lubrication and contribute to disease development.

Here we investigate the role of serine protease CG, with respect to its presence in SF as well as its degrading ability. CG has previously been suggested to be present in SF of OA-joints, using a colorimetric peptide substrate assay, specific for chymotryptic like proteases ^36^. In this study, we validated the finding of CG in OA detecting endogenous CG in SF from 16 patients using western blot (Fig. 2).

A complete degradation of rhPRG4, even for the putatively protected mucin domain, was observed after CG incubation, and this digestion can be enhanced by a prolonged time and increased enzyme levels within physiologically relevant concentrations (Figs 3a, b and Fig. 4a). Moreover, the degrading ability of CG was confirmed introducing CG inhibitor during incubation (Supplementary Fig. S4). These data showed that CG is an efficient protease capable of degrading rhPRG4. Inspired by these results we monitored the effect of CG digestion on native lubricin from SF from individual OA patients and from an RA pool. Similar degrading ability for native lubricin as with recombinant was achieved (Figs 3c, 4a), and a near complete loss of full-length lubricin was observed after incubating with OA SF samples (Figs 4b, 5c). The findings demonstrate the potent lubricin degrading ability of CG, both in a purified *in vitro* incubation system and in a more complex biological environment as in SF form OA patients. The lubricin degrading effect of CG was less pronounced when adding CG to lubricin in SF compared to purified lubricin (Supplementary Fig S5). Degrading efficiency may be influenced by differences in glycosylation of the two native lubricin variants (OA versus RA)^42^, and also for rhPRG4, the latter having CHO cell *O*-glycosylation. We hypothesize that the CG degrading efficiency in SF may also be affected by other SF located proteins competing with lubricin as CG substrates, or by serine protease inhibitors preventing CG activity.

The CG digests of both recombinant and native lubricin revealed a glycosylated lubricin degradation product of approximately 25 kDa, which was detected by western blot (using mAb 9G3 or PNA) or with the glycosensitive stain PAS (Figs 3a, 4a, 4c, and 5c). The presence of glycosylation makes the fragment barely detectable with Coomassie protein stain, tryptic digests of the 25 kDa fragment did not generate any peptides or glycopeptides, which may be expected due to dense and heterogeneous glycosylation, as well as MS ionization inefficiency. However, the data leaves little doubt that the 25 kDa fragment in fact originates from the mucin repeat region, since it could be detected with the monoclonal lubricin antibody 9G3, which binds glycosylated forms of the sequence ‘KEPAPTTT’ ^19^, present multiple times in the mucin domain.

With endogenous CG being present in SF of OA patients, we hypothesized that this protease could generate such a mucin domain containing fragment also *in vivo*. The detection of an endogenously produced 25 kDa lubricin fragment with mAb 9G3 in SF from 13 late-stage OA patients (Fig 5a, b) confirmed that lubricin was indeed degraded partially in OA SF by a CG like protease. In all, our experiments thus supported that CG could be a protease candidate for proteolytic digestion of lubricin in SF. This reflects a local effect of CG during OA pathogenesis, which is in consistent with and extends previous findings that joint inflammation increases during OA and plasma neutrophil activation can serve as a biomarker of OA^44^. Our finding is in accordance with previous knowledge that CG is notably released during inflammatory processes and contributes to ECM degradation^34^. CG as a traditional immune activator and regulator, in a recent research was proved to be responsible for pro-inflammatory IL-1 family cytokines activation^45^. Cytokine IL-1β is found to be active during OA and induces inflammatory responses that causes protease activation and degradation of ECM proteins^46,47^. Furthermore, decreased amount of lubricin was reported by treating cartilage with IL-1β in animal models^28,48^. The present study proves a lubricin degradation ability of CG, which renders CG a potential considerable role during OA, strengthens the importance of further understanding of CG in OA disease and the connection between inflammation and lubricin modifications during the OA pathologic process.

CG has been reported to be a promiscuous protease with both chymase and tryptase activity capable of cleaving both at charged amino acids (eg. Lys and Arg) as well as hydrophobic amino acids (eg Phe, Trp, and Leu)^37,39^. Proteomics and glycoproteomics of a CG digest of recombinant lubricin proved the enzyme to be an efficient protease, and gave an explanation for the degradation of lubricin observed in our western blots. We could detect peptides in the size range of 6-37 amino acids from both the *N*- and *C*-terminal, as well as the mucin domain, proving that CG digested lubricin at numerous sites in the protein backbone. 33 % of the identified peptides contained *C*-terminal Lys residues (Supplementary Table S1 and S2). Hence, the Lys residues which are frequently found throughout the lubricin repeat region in the mucin domain must be obvious targets for digestion. Lys is the most common amino acid (165 aa of total of 1404) in lubricin after Thr and Pro, the two main constituents of the mucin domain. We detected three glycopeptide sequences originating from the mucin domain (Supplementary Table S2). Limited peptide coverage within the mucin domain is expected, since glycosylated peptides from the mucin domain are often more difficult to detect (high heterogeneity, poor ionization efficiency etc). These three glycopeptides occur multiple times in the mucin domain. Together with the detection of two nonglycosylated peptides from the same region (Supplementary Table S1), they provide ample evidence for that the mucin domain, which with glycosylation has an estimated size of 250 kDa (Fig 1), can be degraded into 25-kDa or smaller fragments, as observed with western blot and proteomics.

In order to validate some of the lubricin peptides proposed to be formed by CG digestion in this study, we investigated the presence of ‘in-vitro formed’ autoproteolytic peptides in tryptic digests of recombinant lubricin (rhPRG4). Surprisingly, we did detect a small number of semitryptic peptides, which indicates that larger tryptic lubricin peptides may be degraded in the test tube to smaller semitryptic peptides that mistakenly can be assigned as a protease products. One example was the peptide H.VFMPEVTPDMDYLPR.V (aa1113-1127). The non-tryptic cleavage site at aa 1112 has previously been reported to be a cathepsin S cleavage site in tear fluid ^30^, and equivalent site reported to be an endogenous protease cleavages in SF derived lubricin from horse^44^. Our data suggests an alternative explanation of post-proteolysis induced after tryptic digestion.

OA is a multifactorial disease, and beyond all doubt there is more than one factor that contributes to OA lubricin modifications. The present work demonstrates the potency of CG for lubricin degradation, providing the hypothesis that CG is involved and contributes to OA disease development. The proteolytic cleavage product identified here has a potential to serve as a future biomarker for lubricin degradation in OA.

## Materials and methods

### Lubricin and synovial fluid samples

Synovial fluid (SF) samples were collected from 16 late-stage idiopathic OA patients (8 males and 8 females) subjected to knee replacement surgery. The mean age of the patients was 71 years (range 62-87 years). All individuals gave written consent and the procedure was approved by the ethics committee at Sahlgrenska University Hospital (ethical application 172-15). The SF samples were collected prior surgery, centrifuged, aliquoted and stored at -80°C until assayed. Recombinant lubricin (rhPRG4, 1 mg/mL in Phosphate Buffered Saline (PBS) + 0.1% Tween 20) was obtained from Lubris BioPharma, USA. CG (22 ng/μL in PBS) was isolated from human leukocytes (Sigma-Aldrich, Germany). Purified native lubricin (SF Lub) was enriched from patients’ synovial fluid (pooled from RA patients, n=10) by anion exchange chromatography and ethanol precipitation as described elsewhere^49^, and was quantified using lubricin sandwich ELISA as described below.

### *In vitro* digestion of lubricin by CG

rhPRG4 and lubricin purified from SF, 2-5 μg, were incubated with different concentrations (2.5-125 ng) of CG in at 37°C in PBS (pH 7.4) for 30 minutes or over-night (16 hours). SF (2-2.5 μL) were incubated with CG (22-44 ng) in 37°C for 1 or 2 hours, or a time series of 30 min and 16 hours (specific digestion conditions are indicated in figures).

### SDS-PAGE

Purified native lubricin and SF samples from CG incubations, and also SF samples (16 OA patient samples, 2 μl SF) used for screening of endogenous CG or lubricin degradation fragments, were reduced with 50 mM dithiothreitol (DTT) (Merck KGaA) followed by boiling at 95°C for 15 minutes, and alkylation by 125 mM iodoacetamide (Sigma-Aldrich, St. Louis, MO, US) in dark for 45 minutes. Samples were analysed on NuPAGE Tris-acetate 3-8% gel (Invitrogen, Thermo Fisher Scientific, Waltham, MA, US). One μg of rhPRG4 or 22 ng CG (for endogenous CG assay) were included on each gel as controls. Molecular weight was compared to PageRuler™ Plus Prestained Protein Ladder (10 to 250 kDa, Thermo Fisher Scientific, USA). Gels were stained either with Coomassie brilliant blue R-250 (Bio-Rad Laboratories, Hercules, CA, US) or with Periodic Acid-Schiff (PAS) (Sigma-Aldrich) according to the manufacturers’ instructions. Lubricin band intensities were plotted and peak values were calculated by ImageJ (ImageJ 1.50i, USA)^50^.

### Western blot antibody and lectin staining

After electrophoresis, the gels were blotted to an Immobilon-P PVDF Membrane (Merck Millipore, Burlington, MA, US) using Trans-Blot^®^ SD Semi-Dry Transfer Cell (Bio-Rad Laboratories) at 200 mA for 80 minutes. After blocking with 1% bovine serum albumin (BSA) (VWR, Radnor, PA, US), the membranes were probed with 1 μg/mL mAb 9G3 against the glycosylated epitope ‘KEPAPTTT’ in the lubricin mucin domain (Merck KGaA, Darmstadt, Germany ^51^) or polyclonal anti-CG antibody (Abcam, UK) 1/1000 diluted in assay buffer (1% BSA in PBS-Tween) for detection of endogenous CG in SF, followed by Horseradish Peroxidase (HRP) conjugated goat anti-mouse IgG (H+L) highly cross-adsorbed secondary antibody (Invitrogen, USA) 1/4000 diluted in assay buffer. For the lectin staining assay, blots were first probed by 1 μg/ml biotinylated Peanut Agglutinin (PNA, Vector laboratories, USA) followed by 0.2 μg/ml HRP-streptavidin (Vector laboratories, USA). After incubations, membranes were stained by WesternBright ECL Spray (Advansta, USA) and visualized in a luminescent image analyser (LAS-4000 mini, Fujifilm, Japan). Band intensities were calculated by ImageJ 1.50i^50^.

### Lubricin sandwich ELISA

An in-house ELISA method was set-up and validated for measuring lubricin concentrations in SF. Monoclonal antibody 9G3 (1 μg/mL in PBS) was coated on 96-well Nunc-Immuno maxisorp plates (Thermo Fisher Scientific) at 4°C over-night. After blocking with 3% BSA in PBS+0.05% Tween, SF samples were added as a dilution series (1/50) in assay buffer (1% BSA in PBS-Tween) and incubated for 1 hour at room temperature (RT). Bound proteins were then incubated with biotinylated PNA (Vector laboratories, CA, USA) (1 μg/mL, 1 hour at RT), followed by HRP-streptavidin (Vector laboratories) at 0.1 μg/mL (1 hour at RT). Between each incubation, the wells were washed three times with PBS-Tween to remove unbound reagents. Proteins were stained with 1-Step™ Ultra TMB-ELISA Substrate Solution (Thermo Fisher Scientific) until blue colour was fully generated and the reaction stopped by adding 1 M H_2_SO_4_. Absorbances were read at 450 nm, and compared with a standard curve using recombinant lubricin (dilution series of rhPRG4 (1mg/mL) in assay buffer). Samples were measured in duplicates and mean values are reported. The lubricin ELISA had an intra plate CV = 7.5 % (n=1 SF sample with 10 repeats) and a inter plate CV% =7.8% (n=1 SF sample, tested on 4 plates).

### LC-MS/MS and MS data analyses

rhPRG4 (10 μg in 10 μl PBS + 0.1% Tween 20) was incubated with CG (0.22 μg in 10 μL) in 37°C under non-reducing conditions over night. Incubations were performed with or without subsequent Tween removal (Pierce Detergent Removal Spin Column 125 μL, Thermo Fisher Scientific), but the latter approach generated fewer detected peptides and were therefore not reported here. For control experiments and semitryptic searches, rhPRG4 was digested with trypsin in-solution or in-gel as described elsewhere^52^. The peptides were desalted using C18 ziptips, followed by separation with LC-MS using in-house packed C18 columns at a flow rate of 200 nL/min, and a 45-min gradient of 5-40% buffer B (A: 0.1% formic acid, B:0.1% formic acid, 80% acetonitrile). The column was connected to an Easy-nLC 1000 system (Thermo Fisher Scientific, Odense, Denmark), a nano-electrospray ion source and a Q-Exactive Hybrid Quadrupole-Orbitrap Mass Spectrometer (Thermo Fisher Scientific). For full scan MS, the instrument was scanned *m/z* 350-2000, resolution 60000 (*m/z* 200), AGC target 3e6, max IT 20 ms, dynamic exclusion 10 sec. The twelve most intense peaks (charge states 2, 3, 4) were selected for fragmentation with higher-energy collisional dissociation (HCD). For MS/MS, resolution was set to 15000 (*m/z* 200), AGC target to 5e5, max IT 40 ms, and collision energy NCE=27%.

Raw data files were searched against the human Uniprot protein database (downloaded 17.11.2017) using Peaks Studio 8.5 (Bioinformatics Solutions Inc., Waterloo, Canada). For peptide identification, mass precursor error tolerance was set to 5 ppm, and fragment mass error tolerance to 0.03 Da, enzyme: none; variable modifications: oxidation (M) and deamidation (NQ). Peptide-spectrum matches were filtered to 0.1 % false discovery rate (Peaks peptide score >25). Glycopeptide MS-MS spectra were selected with Peaks Studio software and/or manually using glycan diagnostic ions in the lower mass range (*m/z* 186, 204, 274, 292, 366). Spectra were evaluated manually, all major fragment ions were assigned and > 4 b/y ions required in order to identify the peptide backbone. The assignments were aided by proteomics mining tools available free of charge (Findpept and PeptideMass (web.expasy.org); MS-product (Protein Prospector http://prospector.ucsf.edu). Mass precursor errors were less than 5 ppm.

## Acknowledgements

This study was funded by grants for the Swedish state under the agreement between the Swedish government and the county council, the ALF-agreement (ALFGBG-722391), the Swedish Research Council (621-2013-5895), Kung Gustav V:s 80-års foundation, Petrus and Augusta Hedlund’s foundation (M-2016-0353), AFA insurance research fund (dnr 150150) and IngaBritt and Arne Lundberg Foundation. Sofia Grindberg and Paula-Therese Kelly Pettersson at Danderyd’s Hospital and Lotta Falkendahl at University of Gothenburg are acknowledged for their assistance in collecting samples. Lubris BioPharma, LLC for providing rhPRG4.

## Ethical approval

All OA and RA patients gave informed consent and all the procedures were approved by the regional ethical review board in Gothenburg (172-15,13/5-2015). The study conformed to the ethical guidelines of the 1975 Declaration of Helsinki of research involving human subjects. All methods were performed in accordance with the relevant guidelines and regulations.

## Author contributions

AS and SK suggested the idea and performed initial investigation of lubricin digestion; SH, SA and KAT performed the CG experiments and analysed the data; CJ purified patient lubricin; TAS and GDJ provided rhPRG4; LIB, OR, RK and TE provided patient material. SH, KAT, and NGK wrote the initial manuscript draft; KAT, NGK and TE updated the manuscript. All authors discussed the results, commented, and approved the final manuscript.

