## Supplementary Tables and Figures for "Cathepsin g degrades synovial fluid lubricin: relevance for osteoarthritis pathogenesis"

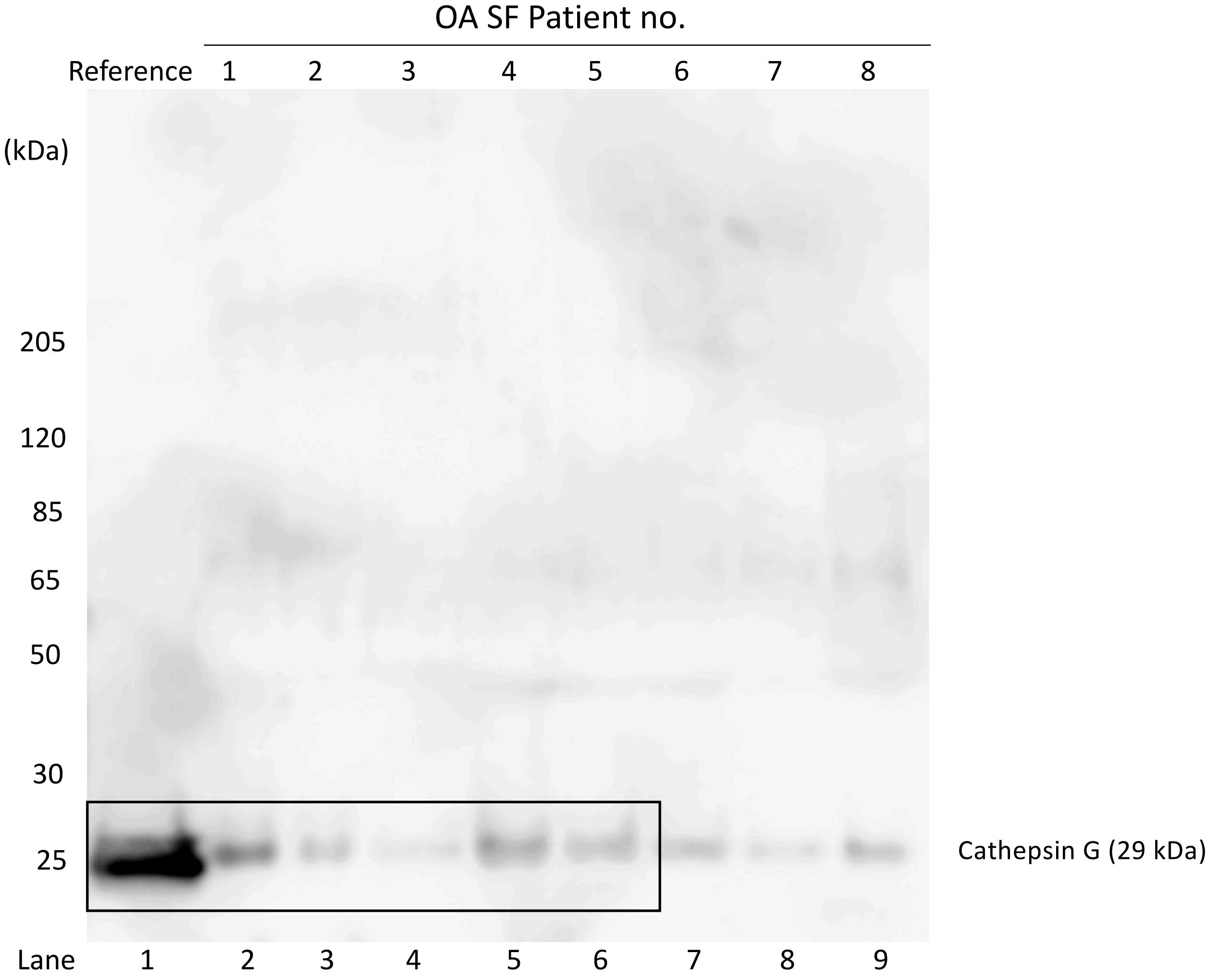

**Supplementary Fig S1**. Endogenous Cathepsin G (CG) in synovial fluid (SF) from 8 OA patients. SF was analysed with SDS-PAGE followed by western blot using a polyclonal anti-CG antibody (see Materials and Methods section). The reference compound was 22 ng CG. The selection of the blot displayed in Figure 2b is marked with a box.

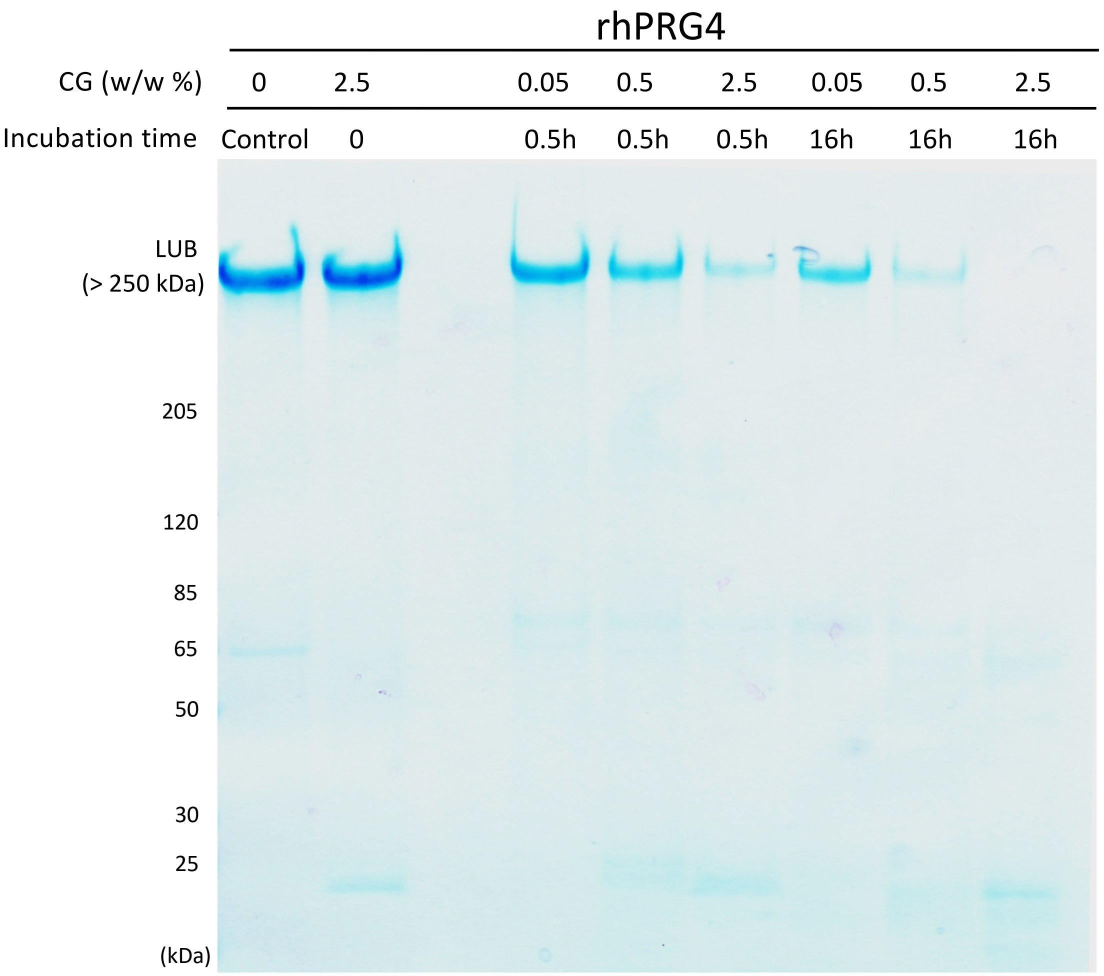

**Supplementary Figure S2. Cathepsin G degradation of recombinant lubricin**. Recombinant lubricin (rhPRG4) (2.5 μg) was incubated with cathepsin G (CG) in PBS at 37°C for 0, 0.5 or 16 hours with different CG/rhPRG4 weight to weight ratios. After reduction and alkylation, the samples were separated on tris-acetate gels (3-8%) and stained with Coomassie Blue (as described in Materials and Methods section).

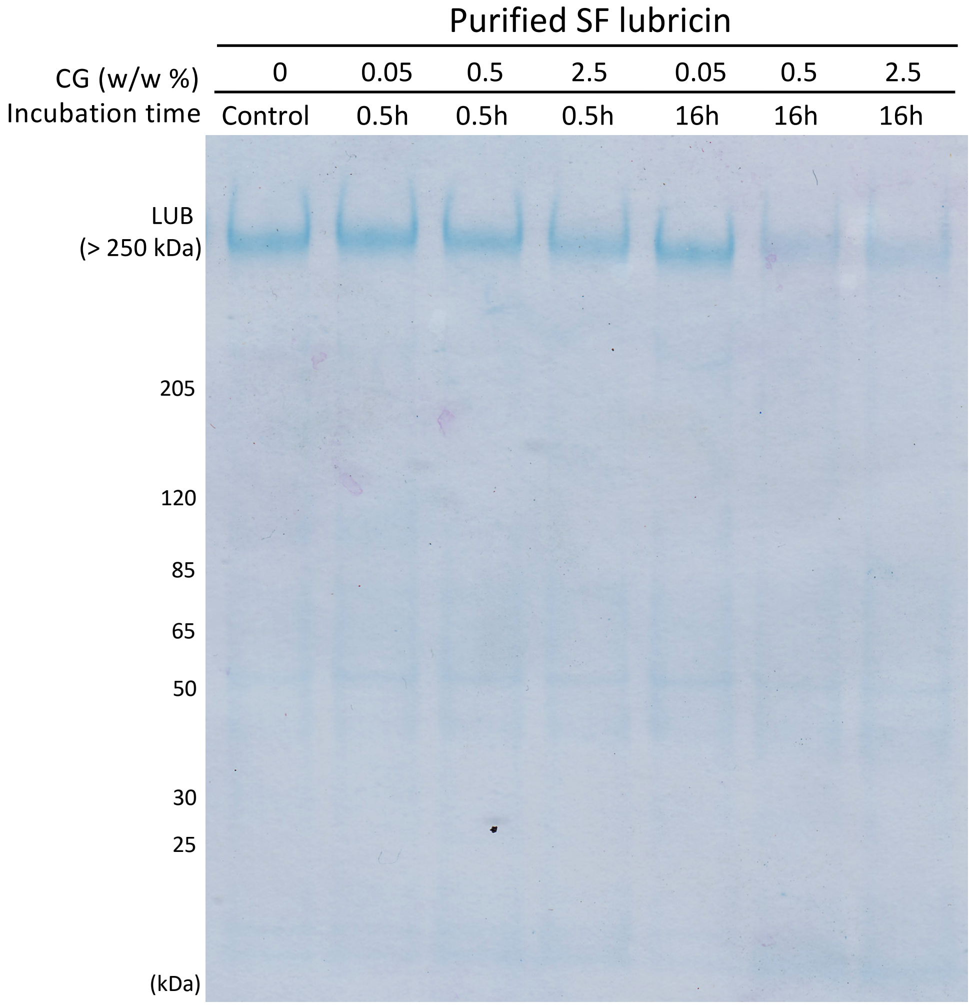

**Supplementary Figure S3. Cathepsin G degradation of lubricin purified from synovial fluid.** Lubricin purified from synovial fluid (2.5 μg) was incubated with cathepsin G (CG) in PBS at 37°C for 0, 0.5 or 16 hours with different CG/rhPRG4 weight to weight ratios. After reduction and alkylation, the samples were separated on tris-acetate gels (3-8%) and stained with Coomassie Blue (as described in Materials and Methods section).

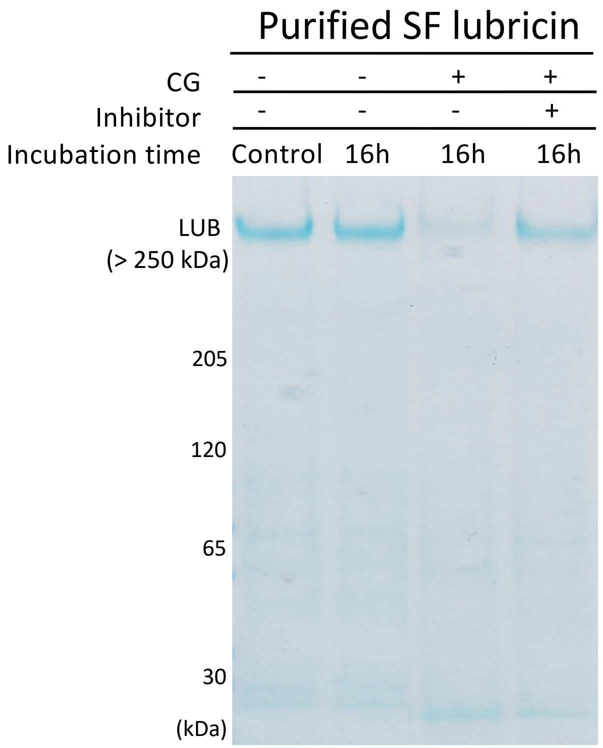

**Supplementary Figure S4. The degradation of lubricin was abolished by the presence of cathepsin G inhibitor.** Purified native lubricin (2.5 μg) from synovial fluid (SF) was incubated with or without cathepsin G (2.5 w/w%) and with or without cathepsin G inhibitor (1 μg, Abcam, UK) in PBS at 37°C over-night for 16 hours. After reduction and alkylation, the samples were separated on tris-acetate gels (3-8%), and stained with Coomassie Blue (as described in Materials and Methods section).

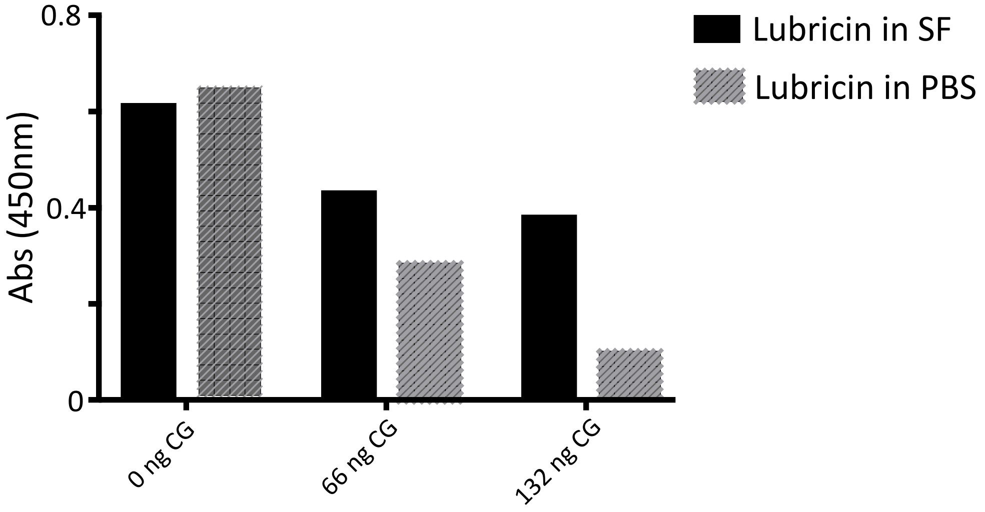

**Supplementary Fig S5**. **Cathepsin G degradation of native lubricin is decreased in synovial fluid compared to in PBS**. SF (5 µl) from an OA patient and purified lubricin from SF (2.5 µg in 5 µl PBS) were incubated with Cathepsin G in PBS (0, 3 and 6 µl, 22ng/ µl) equivalent to 0, 2.5 and 5 w/w%) at 37°C for two hours. Lubricin was detected using a sandwich ELISA, with mAb 9G3 as the catching Ab and PNA lectin for detection (see Materials and Methods section). Samples were measured in duplicates and mean values are shown.

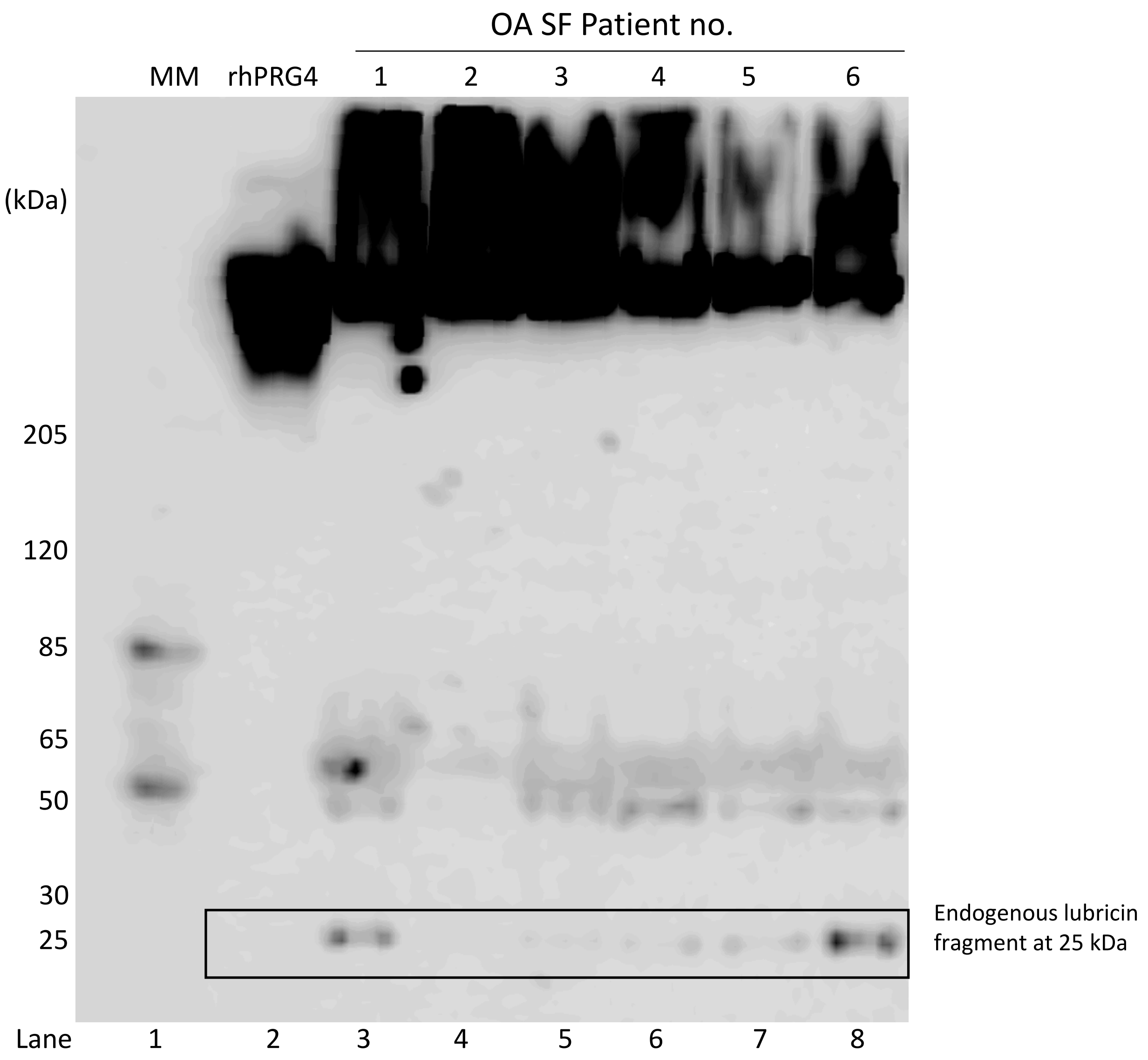

**Supplementary Fig S6**. Identification of the 25 kDa glycosylated lubricin fragment from synovial fluid (SF) of OA patients**.** SF samples (2 µl) were analysed with SDS-PAGE, followed by western blot using mAb 9G3 (as described in Materials and Methods sectionrhPRG4 (1 µg) was used as negative control. ). MM= molecular marker. The selection of the blot displayed in Figure 5a is marked with a box.

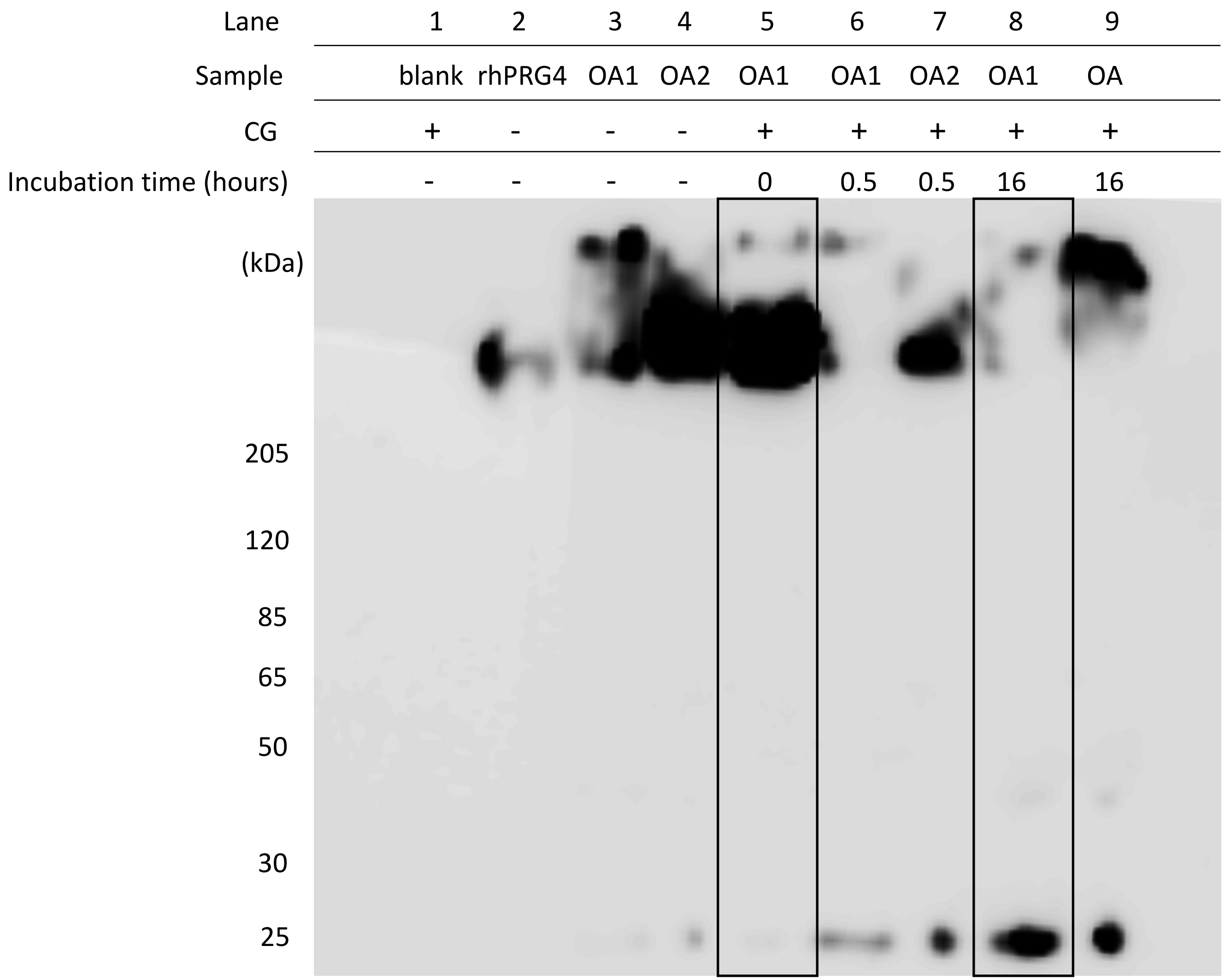

**Supplementary Figure S7.** SF (2 µL) from two OA patients (OA1 and OA2) were incubated with CG (44 ng) for 0, 30 or 16 hours. After reduction and alkylation, the samples were separated on tris-acetate gels (3-8%), and developed with western blot using mAb 9G3 (as described in Materials and Methods section). The selections of the blot displayed in Figure 5c are marked with boxes (lanes 5 and 8).

D.MD**Y**LPRVPN.Q + HexNAc-Hex-NeuAc
Amino acid position in sequence: 1122-1130
Obs. *m/z* 880.8947 (2+)
Calc. $[M+H]^+$ = 1760.7782 ($\Delta$ppm= 2.25)

**Supplementary Figure S8**
HCD spectra of a *O*-glycopeptide (proposed Tyr O-glycosylation) from the mucin domain of recombinant lubricin (rhPRG4) digested with cathepsin G, and analysed with LC/MS/MS using a Qexactive instrument. Analytical conditions are described in Materials and Methods. Diagnostic glycan ions are detected in the lower mass range (*m/z* 100-400). b/y-ions are detected without glycan substituents.

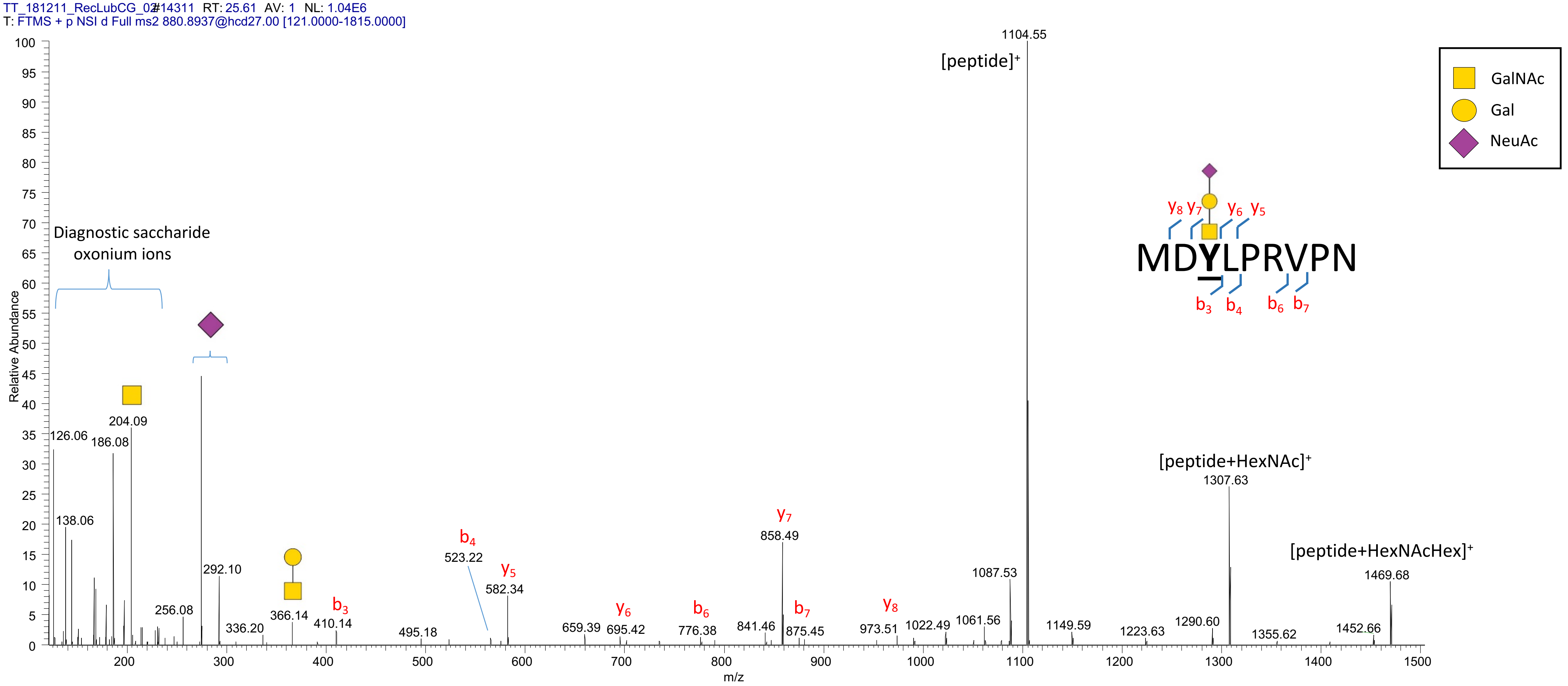

L.R**N**GTVLAF.R+ HexNAc(2) Hex(5)
Amino acid position in sequence: 1158-1165
Obs. *m/z* 1047.4613 (2+)
Calc. $[M+H]^+$ = 2093.9127 ($\Delta$ppm= 1.23 )

**Supplementary Figure S9**

HCD spectra of a *N*-glycopeptide from the mucin domain of recombinant lubricin (rhPRG4) digested with cathepsin G, and analysed with LC/MS/MS using a Qexactive instrument. Analytical conditions are described in Materials and Methods. b/y-ions are detected with or without glycan substituents as annotated.

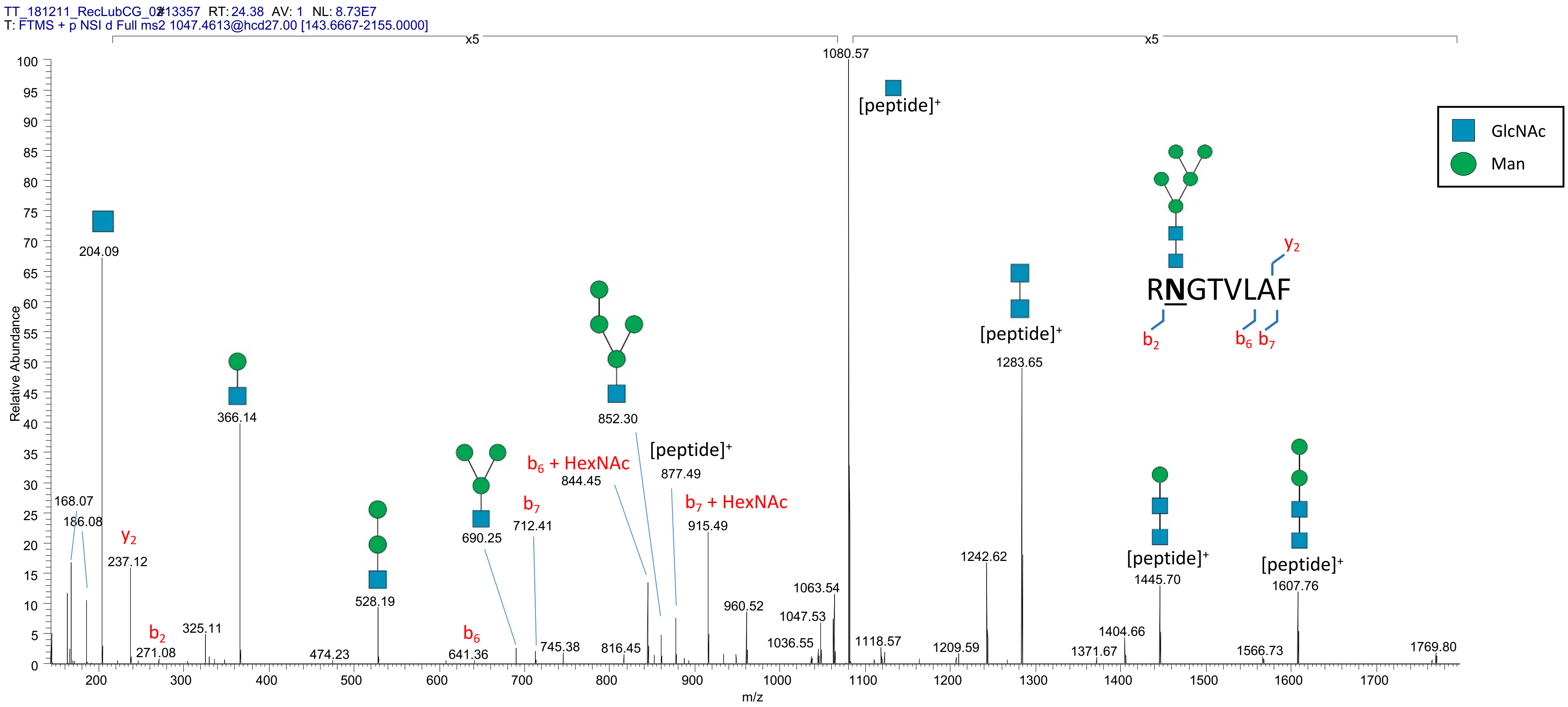

KEPAP**TT**PK+ HexNAc-Hex-NeuAc

Amino acid positions in sequence: 402-10;433-41;449-57;472-80;496-504;558-66;566-74;582-90;606-14;678-86;686-94;694-702;718-26;762-71;771-79;787-95;832-40

Observed *m/z:* 542.2612 (3+)

Calc. $[M+H]^+$ = 1624.7687 ($\Delta$ppm= 0.23)

**Supplementary Figure S10**

HCD spectra of an *O*-glycopeptide from the mucin domain of recombinant lubricin (rhPRG4) digested with cathepsin G, and analysed with LC/MS/MS using a Qexactive instrument. Analytical conditions are described in Materials and Methods. Diagnostic glycan ions are detected in the lower mass range (*m/z* 100-400). b/y-ions are detected without glycan substituents if not annotated elsewise. Potential *O*-glycan sites (Ser/Thr) are underlined and in bold.

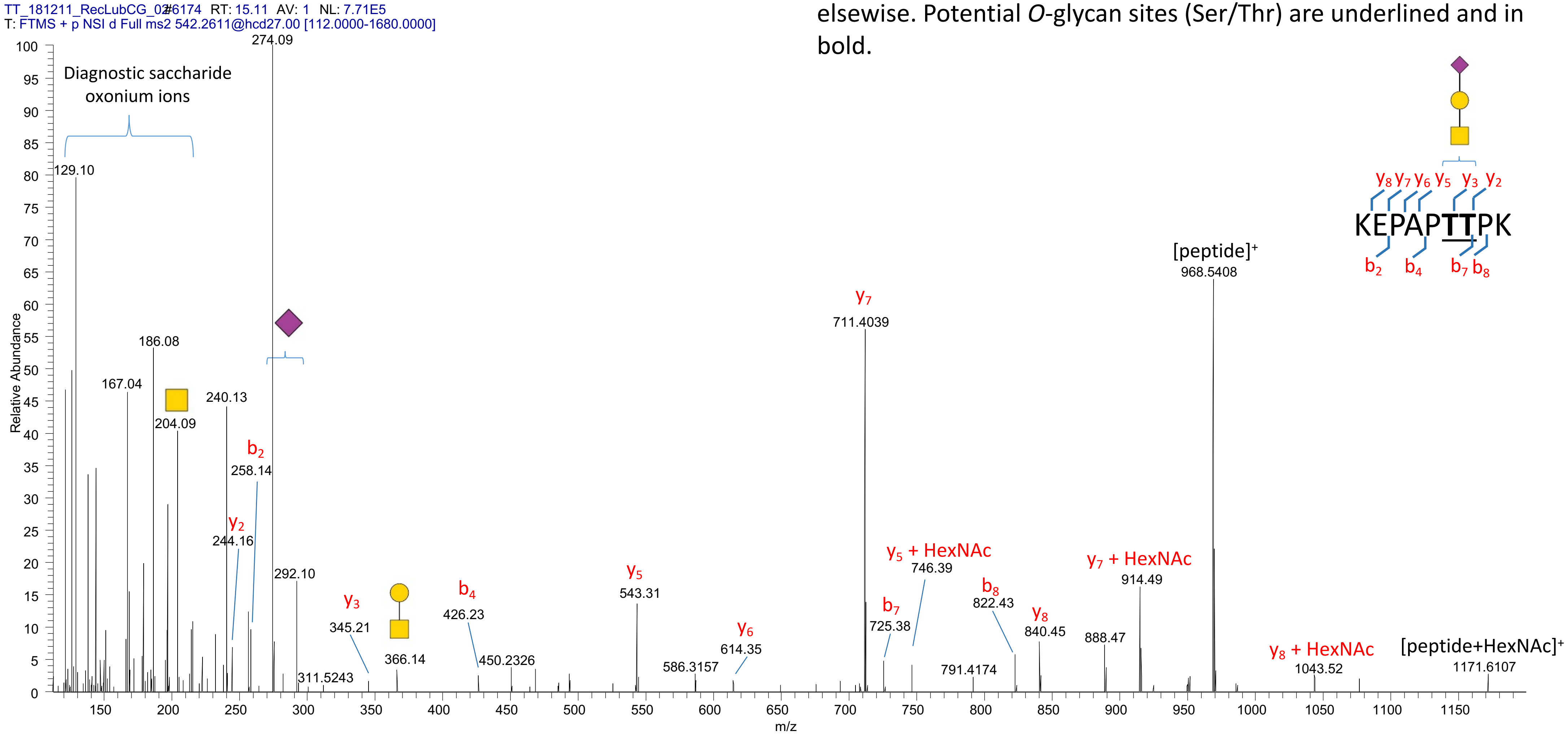

L.KEPAP**TT**PKKPAPK.E + HexNAc(2) Hex(2) NeuAc
(Amino acid position in sequence: 717-731,787-800)
Observed *m/z:* 837.7499 (3+)
Calc. $[M+H]^+$ = 2511.2335 , (Δppm= 0.68)

**Supplementary Figure S11**
HCD spectra of an *O*-glycopeptide from the mucin domain of recombinant lubricin (rhPRG4), digested with cathepsin G, and analysed with LC/MS/MS using a Qexactive instrument. Analytical conditions are described in Materials and Methods. Diagnostic glycan ions are detected in the lower mass range (*m/z* 100-400). b/y-ions are detected without glycan substituents if not annotated elsewise. *O*-glycan sites (Ser/Thr) are underlined and in bold.

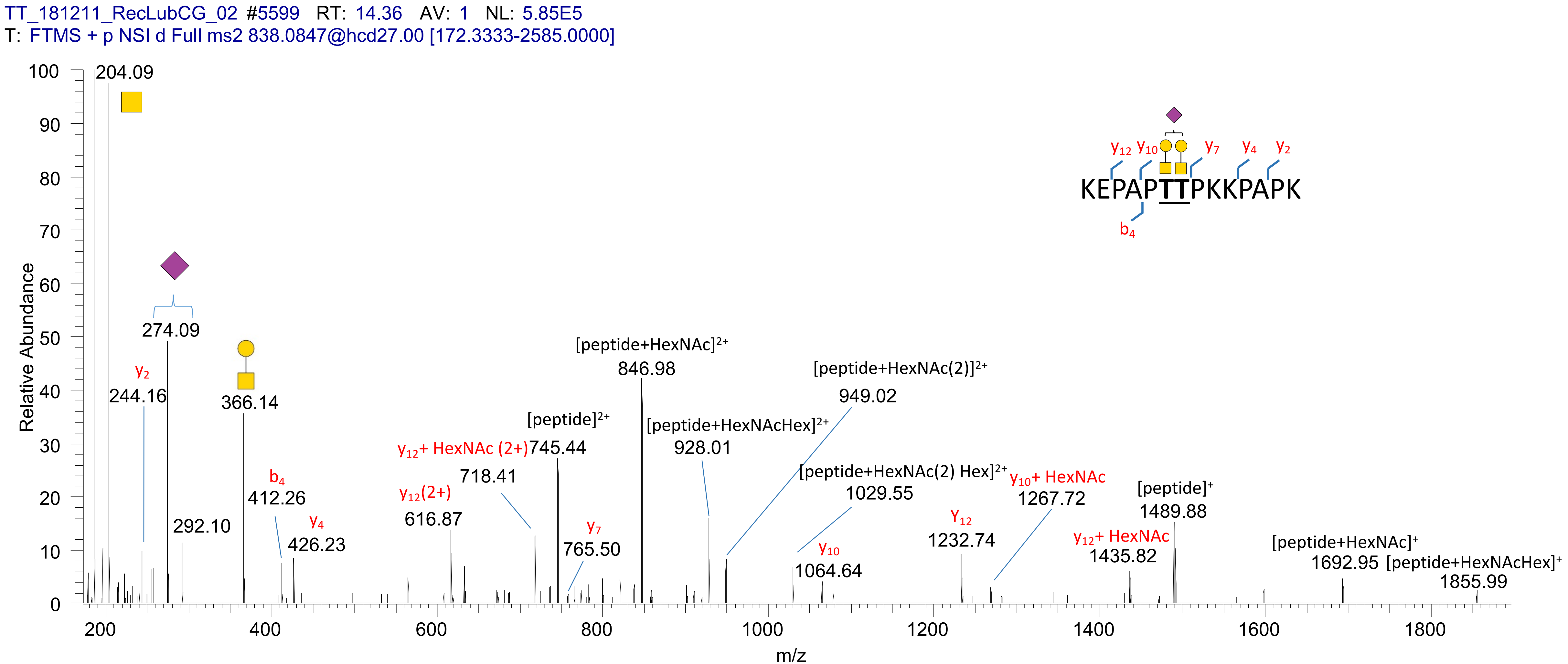

| | | | | | | | | | | | | |
|---|---|---|---|---|---|---|---|---|---|---|---|---|
| **Supplementary Table S1** | | | | | | | | | | | | |
| In-solution cathepsin G digest of recombinant lubricin (rhPRG4) | | | | | | | | | | | | |
| Peptides were analyzed with LC-MS/MS as described in Materials and Methods section | | | | | | | | | | | | |
| Data was searched against the Uniprot database and filtered to 0.1 % FDR with 'no enzyme specificity' settings using Peaks Studio 8.5 as described in Materials and Methods section. | | | | | | | | | | | | |
| Peptides which are nonspecific for the cathepsin G digest are labelled with an asterisk (*). These were also detected in semitryptic searches of tryptic digests of rhPRG4 (data not shown) | | | | | | | | | | | | |
| **Peptide** | **Start** | **End** | **-10lgP** | **Mass** | **ppm** | **m/z** | **z** | **RT** | **Area** | **Scan** | **#Spec** | **PTM** |
| K.KAPPPSGASQTIK.S | 118 | 130 | 44.83 | 1280.709 | 1.5 | 427.9108 | 3 | 14.83 | 5.71E+08 | 5951 | 21 | |
| K.STTKRSPKPPNKK.K | 131 | 143 | 26.52 | 1467.852 | 0.5 | 367.9705 | 4 | 10.93 | 1.88E+07 | 3122 | 2 | |
| K.RSPKPPNKKKTK.K | 135 | 146 | 42.52 | 1407.867 | 0.9 | 352.9744 | 4 | 10.62 | 2.37E+08 | 2999 | 20 | |
| K.RSPKPPNKKKTKK.V | 135 | 147 | 41.99 | 1535.962 | 0 | 384.9979 | 4 | 11.06 | 3.40E+07 | 3206 | 4 | |
| K.RSPKPPNKK.K | 135 | 143 | 33.45 | 1050.63 | 0.5 | 351.2174 | 3 | 5.35 | 1.38E+08 | 1523 | 21 | |
| K.TKKVIESEEITEEH.S | 145 | 158 | 29.82 | 1670.836 | 0.7 | 557.9531 | 3 | 18.31 | 1.61E+07 | 8632 | 2 | |
| K.KVIESEEITEEH.S | 147 | 158 | 39.44 | 1441.694 | 0.8 | 721.8547 | 2 | 19.64 | 1.06E+09 | 9663 | 12 | |
| K.KVIESEEIT.E | 147 | 155 | 25.49 | 1046.55 | 0.1 | 524.2821 | 2 | 20.25 | 6.99E+07 | 10142 | 2 | |
| K.VIESEEITEEH.S* | 148 | 158 | 33.81 | 1313.599 | 1.1 | 657.8073 | 2 | 21.62 | 5.51E+08 | 11206 | 8 | |
| H.SVSENQESSSSSSSSSSSSTIRK.I* | 159 | 181 | 27.61 | 2334.042 | 1.2 | 779.0222 | 3 | 14.61 | 5.02E+06 | 5794 | 1 | |
| S.SSSSSSSSSSSTIRK.I | 167 | 181 | 29.09 | 1473.691 | 0 | 492.2375 | 3 | 12.21 | 4.98E+06 | 3946 | 2 | |
| S.SSSSSSSSSSTIRKIK.S | 168 | 183 | 38 | 1627.838 | -0.1 | 543.6198 | 3 | 14.81 | 2.64E+07 | 5939 | 2 | |
| S.SSSSSSSSSSTIRK.I* | 168 | 181 | 33.67 | 1386.659 | 0.1 | 463.2269 | 3 | 12.32 | 1.11E+07 | 4033 | 2 | |
| S.SSSSSSSSSTIRKIK.S | 169 | 183 | 36.56 | 1540.806 | 1.1 | 514.6097 | 3 | 14.65 | 4.21E+07 | 5820 | 1 | |
| S.SSSSSSSSSTIRK.I* | 169 | 181 | 31.58 | 1299.627 | 0.5 | 434.2163 | 3 | 12.21 | 2.79E+07 | 3942 | 3 | |
| S.SSSSSSSSTIRKIK.S | 170 | 183 | 33.8 | 1453.774 | 0.4 | 485.5987 | 3 | 14.48 | 5.25E+07 | 5689 | 4 | |
| S.SSSSSSSSTIRK.I* | 170 | 181 | 28.01 | 1212.595 | 0.2 | 405.2056 | 3 | 12.09 | 2.39E+07 | 3851 | 2 | |
| S.SSSSSSSTIRKIK.S | 171 | 183 | 31.53 | 1366.742 | 0.3 | 456.588 | 3 | 14.14 | 3.79E+07 | 5429 | 2 | |
| S.SSSSSSSTIRK.I* | 171 | 181 | 27.12 | 1125.563 | 0 | 376.1948 | 3 | 11.86 | 1.40E+07 | 3674 | 2 | |
| S.SSSSSSTIRKIK.S | 172 | 183 | 31.45 | 1279.71 | -0.6 | 427.5769 | 3 | 14.03 | 3.55E+07 | 5344 | 3 | |
| S.SSSSSTIRKIK.S | 173 | 183 | 33.3 | 1192.678 | -0.1 | 597.3459 | 2 | 13.59 | 3.48E+07 | 5015 | 2 | |
| S.SSSSSTIRK.I* | 173 | 181 | 26.29 | 951.4985 | -3.3 | 476.755 | 2 | 11.13 | 4.17E+06 | 3244 | 2 | |
| S.SSSSTIRKIK.S | 174 | 183 | 29.17 | 1105.646 | 0 | 369.5558 | 3 | 13.34 | 2.62E+07 | 4819 | 2 | |
| K.SSKNSAANRELQK.K | 184 | 196 | 35.13 | 1431.743 | -0.5 | 478.2547 | 3 | 12.46 | 5.74E+07 | 4141 | 2 | |
| K.SSKN(+.98)SAANRELQK.K | 184 | 196 | 30.48 | 1432.727 | 0.1 | 478.5829 | 3 | 13 | 7.22E+06 | 4555 | 1 | Deamidatio |
| K.NSAANRELQK.K | 187 | 196 | 27.22 | 1129.584 | 0.2 | 565.7994 | 2 | 13.22 | 2.98E+07 | 4728 | 4 | |
| N.SAANRELQK.K | 188 | 196 | 28.6 | 1015.541 | 0.5 | 508.778 | 2 | 13.06 | 1.02E+08 | 4597 | 2 | |
| F.KVTTPDTSTTQHNK.V | 232 | 245 | 46.16 | 1556.779 | 0.1 | 519.9338 | 3 | 13.14 | 2.72E+07 | 4660 | 3 | |
| F.KVTTPDTSTTQHNKVSTSPK.I | 232 | 251 | 45.7 | 2156.107 | -0.2 | 719.7096 | 3 | 15 | 1.79E+07 | 6086 | 2 | |
| F.KVTTPDTSTTQH.N | 232 | 243 | 37.56 | 1314.642 | 0.6 | 439.2214 | 3 | 14.61 | 1.72E+07 | 5789 | 4 | |
| F.KVTTPDTSTTQHN.K | 232 | 244 | 35.78 | 1428.684 | 0.6 | 477.2357 | 3 | 14.5 | 4.96E+07 | 5708 | 2 | |
| F.KVTTPDTSTTQHN(+.98)KVSTSPK.I | 232 | 251 | 33.79 | 2157.091 | -1.1 | 432.425 | 5 | 15.28 | 3.06E+06 | 6298 | 2 | Deamidatio |
| F.KVTTPDTSTTQ.H | 232 | 242 | 31.81 | 1177.583 | 0.4 | 589.7988 | 2 | 15.42 | 3.22E+07 | 6407 | 2 | |
| K.VTTPDTSTTQHNKVSTSPK.I | 233 | 251 | 43.55 | 2028.012 | 0.6 | 677.0118 | 3 | 15.67 | 2.44E+07 | 6598 | 1 | |
| K.VTTPDTSTTQHNK.V | 233 | 245 | 37.24 | 1428.684 | 0.7 | 477.2357 | 3 | 14.16 | 4.96E+07 | 5442 | 3 | |
| K.VTTPDTSTTQH.N | 233 | 243 | 32.39 | 1186.547 | 0.5 | 594.2809 | 2 | 15.76 | 2.54E+07 | 6663 | 2 | |
| K.VTTPDTSTTQHN.K | 233 | 244 | 31.34 | 1300.59 | -1.4 | 651.3011 | 2 | 15.44 | 3.59E+07 | 6419 | 2 | |
| S.KETSLTVN.K | 272 | 279 | 25.18 | 890.4709 | 0.5 | 446.243 | 2 | 17.57 | 5.86E+06 | 8065 | 2 | |
| L.TVNKETTVETK.E | 277 | 287 | 38.78 | 1248.656 | 1 | 625.3359 | 2 | 13.57 | 8.06E+07 | 5000 | 4 | |
| N.KETTVETK.E | 280 | 287 | 25.1 | 934.4971 | 0.6 | 468.2561 | 2 | 11.27 | 7.22E+07 | 3322 | 3 | |
| N.KQTSTDGKEKTTSAKETQ.S | 294 | 311 | 46.92 | 1966.981 | 1 | 656.6682 | 3 | 11.22 | 1.60E+08 | 3295 | 2 | |
| N.KQTSTDGKEKTTSAK.E | 294 | 308 | 44.36 | 1608.832 | 0 | 403.2152 | 4 | 2.52 | 2.91E+07 | 719 | 1 | |
| N.KQTSTDGKEKTTSA.K | 294 | 307 | 42.14 | 1480.737 | 0.5 | 494.5865 | 3 | 11.01 | 1.42E+07 | 3175 | 1 | |
| N.KQTSTDGKEKTTSAKETQ(+.98)SIEKTSAK.D | 294 | 319 | 31.81 | 2812.43 | 3.8 | 469.7474 | 6 | 13.41 | 7.27E+06 | 4873 | 1 | Deamidatio |
| S.TDGKEKTTSAKETQSIEK.T | 298 | 315 | 46.52 | 1980.001 | 0.2 | 496.0076 | 4 | 13.37 | 2.30E+07 | 4844 | 2 | |
| K.TTSAKETQSIEKT.S | 304 | 316 | 43.51 | 1422.72 | 0.8 | 712.368 | 2 | 15.21 | 5.92E+07 | 6247 | 4 | |
| K.TTSAKETQSIEK.T | 304 | 315 | 41.92 | 1321.673 | 1.7 | 661.8447 | 2 | 14.33 | 2.23E+08 | 5575 | 8 | |
| T.TSAKETQSIEK.T | 305 | 315 | 31.26 | 1220.625 | -0.1 | 407.8821 | 3 | 13.56 | 5.16E+06 | 4989 | 1 | |
| T.SAKETQSIEKTSAK.D | 306 | 319 | 37.68 | 1506.789 | 1.2 | 503.2709 | 3 | 12.95 | 5.86E+06 | 4513 | 2 | |
| A.KETQSIEKTSAK.D | 308 | 319 | 37.52 | 1348.72 | -0.2 | 450.5804 | 3 | 12.31 | 6.21E+06 | 4023 | 2 | |
| K.ETQSIEKTSAK.D | 309 | 319 | 26.09 | 1220.625 | -0.1 | 407.8821 | 3 | 13.13 | 1.10E+07 | 4652 | 2 | |
| K.SAPTTPKEPAPTTTK.S | 373 | 387 | 25.56 | 1525.799 | -1.9 | 509.6059 | 3 | 16.38 | 4.70E+06 | 7139 | 2 | |
| K.SAPTTTKEPAPTTTK.S | 529 | 543 | 25.16 | 1529.794 | -0.1 | 510.9384 | 3 | 14.88 | 2.16E+06 | 5993 | 1 | |
| L.KEPAPTTPKKPAPK.E | 718 | 731 | 46.35 | 1488.866 | -0.3 | 373.2238 | 4 | 12.85 | 4.84E+07 | 4436 | 3 | |
| H.KSPDESTPELS.A | 871 | 881 | 26.37 | 1188.551 | 1 | 595.2834 | 2 | 19.1 | 3.82E+07 | 9243 | 2 | |
| H.KSPDESTPEL.S | 871 | 880 | 25.72 | 1101.519 | 1.4 | 551.7675 | 2 | 21.39 | 1.17E+08 | 11022 | 2 | |
| K.SEDAGGAEGETPHM.L* | 1094 | 1107 | 39.1 | 1386.536 | 1 | 694.2759 | 2 | 19.92 | 2.30E+07 | 9881 | 4 | |
| H.MLLRPHVF.M | 1107 | 1114 | 35.7 | 1011.569 | 0.2 | 506.7917 | 2 | 26.84 | 1.26E+09 | 15249 | 3 | |
| H.M(+15.99)LLRPHVF.M | 1107 | 1114 | 29.98 | 1027.564 | 0.2 | 514.7892 | 2 | 25.21 | 2.69E+08 | 14003 | 3 | Oxidation (I |
| H.MLLRPHV.F | 1107 | 1113 | 29.29 | 864.5004 | -0.7 | 433.2572 | 2 | 22.45 | 5.33E+07 | 11857 | 2 | |
| H.M(+15.99)LLRPHV.F | 1107 | 1113 | 26.94 | 880.4953 | 0.9 | 441.2553 | 2 | 20.67 | 5.03E+06 | 10467 | 2 | Oxidation (I |
| M.LLRPHVF.M | 1108 | 1114 | 28.29 | 880.5283 | -1.9 | 441.2706 | 2 | 24.33 | 4.34E+08 | 13318 | 7 | |
| F.MPEVTPDM.D | 1115 | 1122 | 30.68 | 918.3827 | 1 | 460.1991 | 2 | 26.42 | 4.13E+06 | 14930 | 2 | |
| T.PDMDYLPRVPN.Q | 1120 | 1130 | 38.88 | 1315.623 | 1.1 | 658.8195 | 2 | 28.14 | 4.59E+06 | 16213 | 2 | |
| D.MDYLPRVPNQGI.I | 1122 | 1133 | 46.68 | 1401.707 | 0 | 701.861 | 2 | 29.17 | 3.77E+07 | 16909 | 3 | |
| D.MDYLPRVPN.Q | 1122 | 1130 | 40.69 | 1103.543 | -0.1 | 552.7789 | 2 | 28.1 | 2.16E+09 | 16181 | 10 | |
| M.DYLPRVPN.Q | 1123 | 1130 | 31.73 | 972.5029 | 0.3 | 487.2589 | 2 | 25.82 | 2.49E+08 | 14475 | 9 | |
| N.GKPVDGLTTLR.N* | 1148 | 1158 | 25.66 | 1155.661 | 0.7 | 578.8383 | 2 | 21.95 | 8.78E+06 | 11459 | 1 | |
| W.MLSPFSPPSPARR.I | 1172 | 1184 | 34.74 | 1441.75 | 1.2 | 721.8831 | 2 | 25.82 | 1.38E+08 | 14476 | 6 | |
| W.MLSPFSPPSPAR.R* | 1172 | 1183 | 30.28 | 1285.649 | 1.7 | 643.8328 | 2 | 28.02 | 2.43E+07 | 16120 | 3 | |
| M.LSPFSPPSPARR.I | 1173 | 1184 | 30.2 | 1310.71 | 3.1 | 656.3641 | 2 | 23.05 | 3.59E+08 | 12318 | 9 | |
| M.LSPFSPPSPAR.R* | 1173 | 1183 | 29.56 | 1154.608 | -0.1 | 578.3114 | 2 | 25.12 | 8.46E+07 | 13932 | 4 | |
| F.SPPSPARRITEVW.G | 1177 | 1189 | 33.92 | 1494.794 | 0.3 | 748.4047 | 2 | 26.03 | 2.48E+09 | 14644 | 12 | |
| F.SPPSPARR.I | 1177 | 1184 | 25.78 | 866.4722 | -0.2 | 434.2433 | 2 | 12.56 | 1.13E+08 | 4215 | 5 | |
| E.VWGIPSPIDTVF.T | 1188 | 1199 | 27.19 | 1329.697 | 1.4 | 665.8566 | 2 | 40.5 | 8.76E+06 | 25480 | 1 | |
| V.WGIPSPIDTVF.T | 1189 | 1199 | 26.24 | 1230.628 | 1.7 | 616.3225 | 2 | 39.57 | 5.27E+06 | 24826 | 1 | |
| F.KDSQYWRF.T* | 1212 | 1219 | 25.59 | 1128.535 | -0.3 | 377.1855 | 3 | 26.57 | 1.65E+07 | 15042 | 1 | |
| W.RFTNDIKDAGYPKPIFK.G | 1218 | 1234 | 26.95 | 2009.073 | -0.1 | 402.8219 | 5 | 24.22 | 6.87E+06 | 13227 | 1 | |
| F.TNDIKDAGYPKPIFK.G | 1220 | 1234 | 47.19 | 1705.904 | 4.6 | 569.6445 | 3 | 23.25 | 1.82E+09 | 12479 | 7 | |
| F.TN(+.98)DIKDAGYPKPIFK.G | 1220 | 1234 | 36.53 | 1706.888 | 3.2 | 569.9717 | 3 | 24.17 | 1.80E+07 | 13186 | 1 | Deamidatio |
| F.TNDIKDAGYPKPI.F | 1220 | 1232 | 33.51 | 1430.741 | -0.8 | 477.9204 | 3 | 22.42 | 1.49E+07 | 11832 | 1 | |
| F.TNDIKDAGYPKPIFKGF.G | 1220 | 1236 | 30.59 | 1909.994 | -2.5 | 637.6703 | 3 | 27.65 | 1.47E+07 | 15842 | 1 | |
| T.NDIKDAGYPKPIFK.G | 1221 | 1234 | 44.5 | 1604.856 | 0.8 | 535.9598 | 3 | 23.22 | 1.59E+08 | 12453 | 3 | |
| T.NDIKDAGYPKPIF.K | 1221 | 1233 | 35.73 | 1476.761 | 0.6 | 493.2613 | 3 | 27.18 | 7.83E+06 | 15487 | 1 | |

| | | | | | | | | | | | | |
|---|---|---|---|---|---|---|---|---|---|---|---|---|
| N.DIKDAGYPKPIFK.G | 1222 | 1234 | 40.79 | 1490.813 | 0.3 | 746.4141 | 2 | 24.07 | 1.20E+08 | 13116 | 6 | |
| N.DIKDAGYPKPIF.K | 1222 | 1233 | 35.5 | 1362.718 | 0.9 | 682.367 | 2 | 28.15 | 1.12E+07 | 16224 | 2 | |
| I.KDAGYPKPIFK.G | 1224 | 1234 | 37.46 | 1262.702 | -0.1 | 421.908 | 3 | 21.09 | 1.16E+07 | 10788 | 2 | |
| K.DAGYPKPIFK.G | 1225 | 1234 | 38.12 | 1134.607 | 2.8 | 568.3125 | 2 | 23.4 | 6.77E+09 | 12593 | 20 | |
| K.DAGYPKPIFKGF.G | 1225 | 1236 | 36.03 | 1338.697 | 2 | 670.3572 | 2 | 29.22 | 6.68E+06 | 16937 | 1 | |
| K.DAGYPKPI.F | 1225 | 1232 | 25.17 | 859.4439 | -0.3 | 430.7291 | 2 | 22.17 | 3.24E+07 | 11631 | 2 | |
| D.AGYPKPIFK.G | 1226 | 1234 | 25.38 | 1019.58 | -2.2 | 510.7963 | 2 | 23.42 | 3.55E+06 | 12610 | 2 | |
| A.GYPKPIFK.G* | 1227 | 1234 | 29.13 | 948.5432 | -0.9 | 475.2784 | 2 | 23.48 | 2.39E+07 | 12657 | 3 | |
| K.GFGGLTGQIVAAL.S | 1235 | 1247 | 28.51 | 1202.666 | 1.1 | 602.3409 | 2 | 37.1 | 1.80E+08 | 22941 | 3 | |
| K.GFGGLTGQIV.A | 1235 | 1244 | 25.07 | 947.5076 | -0.3 | 474.761 | 2 | 31.9 | 3.98E+07 | 18736 | 1 | |
| Y.KNWPESVYF.F | 1253 | 1261 | 26.08 | 1168.555 | 0.1 | 585.285 | 2 | 30.78 | 2.16E+08 | 17967 | 1 | |
| Y.FFKRGGSIQQY.I | 1261 | 1271 | 28.63 | 1329.683 | -0.1 | 444.2348 | 3 | 21.5 | 3.94E+07 | 11113 | 2 | |
| F.FKRGGSIQQY.I | 1262 | 1271 | 27.29 | 1182.615 | 1.2 | 592.3152 | 2 | 18.5 | 3.13E+08 | 8786 | 6 | |
| L.NYPVYGETTQVR.R | 1289 | 1300 | 29.13 | 1425.689 | -3.7 | 713.8491 | 2 | 23.46 | 9.15E+06 | 12640 | 2 | |
| R.FERAIGPSQTH.T | 1304 | 1314 | 28.71 | 1241.615 | -1.5 | 414.8784 | 3 | 17.94 | 2.12E+07 | 8349 | 3 | |
| F.ERAIGPSQTH.T | 1305 | 1314 | 32.6 | 1094.547 | 0.7 | 365.8565 | 3 | 14.77 | 6.36E+05 | 5906 | 1 | |
| R.AIGPSQTHTIR.I | 1307 | 1317 | 39.4 | 1179.636 | -1.9 | 394.2185 | 3 | 17.27 | 1.32E+08 | 7830 | 5 | |
| R.AIGPSQTHTI.R | 1307 | 1316 | 30.69 | 1023.535 | 1.2 | 512.7753 | 2 | 20.92 | 1.25E+07 | 10660 | 2 | |
| R.AIGPSQTH.T | 1307 | 1314 | 29.42 | 809.4031 | -0.6 | 405.7086 | 2 | 15.64 | 3.22E+08 | 6575 | 8 | |
| I.GPSQTHTIR.I | 1309 | 1317 | 30.2 | 995.5148 | 1.4 | 498.7654 | 2 | 17.44 | 8.13E+05 | 7961 | 1 | |
| T.HTIRIQYSPARL.A | 1314 | 1325 | 26.91 | 1453.815 | 0.9 | 485.6128 | 3 | 23.33 | 2.01E+06 | 12541 | 1 | |
| H.TIRIQYSPARL.A | 1315 | 1325 | 36.27 | 1316.757 | 0.9 | 439.9265 | 3 | 25.55 | 8.91E+08 | 14268 | 7 | |
| H.TIRIQYSPAR.L | 1315 | 1324 | 36 | 1203.672 | -1.4 | 402.2308 | 3 | 21.61 | 1.21E+07 | 11192 | 2 | |
| H.TIRIQYSPARLA.Y | 1315 | 1326 | 34.7 | 1387.794 | -1.1 | 463.6046 | 3 | 25.23 | 1.57E+06 | 14017 | 1 | |
| T.IRIQYSPARL.A | 1316 | 1325 | 29.52 | 1215.709 | 0 | 406.2435 | 3 | 25.01 | 3.08E+06 | 13843 | 1 | |
| I.RIQYSPARL.A | 1317 | 1325 | 26.79 | 1102.625 | -0.6 | 368.5486 | 3 | 22.11 | 3.67E+07 | 11589 | 2 | |
| R.IQYSPARL.A | 1318 | 1325 | 25.51 | 946.5236 | -0.2 | 474.269 | 2 | 24.02 | 2.16E+08 | 13076 | 1 | |
| L.AYQDKGVLH.N | 1326 | 1334 | 31.51 | 1029.524 | 0.8 | 515.7698 | 2 | 17.19 | 1.03E+07 | 7766 | 2 | |
| A.YQDKGVLH.N | 1327 | 1334 | 26.49 | 958.4872 | 0.6 | 480.2512 | 2 | 16.72 | 2.97E+06 | 7399 | 1 | |
| Y.QDKGVLH.N | 1328 | 1334 | 25.24 | 795.4239 | -0.2 | 398.7191 | 2 | 13.39 | 9.53E+07 | 4856 | 2 | |
| K.VSILWRGLPNVVTSAISLPN(+.98)IRKPDGYDYYAFSKDQYY.N | 1339 | 1376 | 32.53 | 4409.237 | 4.6 | 1103.322 | 4 | 35.85 | 4.26E+06 | 21950 | 1 | Deamidatio |
| L.WRGLPN.V | 1343 | 1348 | 30.55 | 741.3922 | 0.9 | 371.7037 | 2 | 23.38 | 1.50E+09 | 12578 | 9 | |
| L.WRGLPNVVTS.A | 1343 | 1352 | 27.63 | 1127.609 | 0.6 | 564.812 | 2 | 27.74 | 7.66E+08 | 15906 | 5 | |
| W.RGLPNVVTSAI.S | 1344 | 1354 | 34.26 | 1125.651 | 0 | 563.8326 | 2 | 28.39 | 3.77E+08 | 16383 | 4 | |
| N.VVTSAISLPNIRKPDGY.D | 1349 | 1365 | 28.26 | 1829.005 | 1 | 610.6761 | 3 | 27.9 | 3.05E+07 | 16027 | 1 | |
| V.TSAISLPNIRKPDGY.D | 1351 | 1365 | 40.62 | 1630.868 | 1.1 | 816.4421 | 2 | 26.63 | 1.24E+08 | 15091 | 4 | |
| T.SAISLPNIRKPDGY.D | 1352 | 1365 | 37.49 | 1529.82 | 1.3 | 510.948 | 3 | 26.7 | 3.13E+09 | 15144 | 12 | |
| T.SAISLPNIRK.P | 1352 | 1361 | 25.44 | 1097.656 | -0.8 | 549.8347 | 2 | 24.01 | 2.74E+07 | 13064 | 3 | |
| S.AISLPN(+.98)IRKPDGY.D | 1353 | 1365 | 29.23 | 1443.772 | 0.2 | 482.2647 | 3 | 27.36 | 2.04E+07 | 15624 | 1 | Deamidatio |
| S.AISLPNIRKPDGYDYY.A | 1353 | 1368 | 28.3 | 1883.942 | 0.1 | 942.9781 | 2 | 29.01 | 8.35E+08 | 16809 | 3 | |
| A.ISLPNIRKPDGY.D | 1354 | 1365 | 32.61 | 1371.751 | -0.3 | 458.2575 | 3 | 26.19 | 8.01E+07 | 14756 | 6 | |
| I.SLPNIRKPDGYDYY.A | 1355 | 1368 | 43.43 | 1699.821 | 1.9 | 850.9192 | 2 | 25.93 | 2.80E+08 | 14564 | 3 | |
| L.PNIRKPDGY.D | 1357 | 1365 | 38.04 | 1058.551 | -0.3 | 530.2826 | 2 | 22.79 | 1.65E+07 | 12116 | 2 | |
| N.IRKPDGY.D | 1359 | 1365 | 26.16 | 847.4551 | -0.2 | 424.7348 | 2 | 16.62 | 9.86E+07 | 7321 | 3 | |
| Y.AFSKDQYY.N | 1369 | 1376 | 31.31 | 1020.455 | 1.4 | 511.2356 | 2 | 22.37 | 1.59E+09 | 11793 | 6 | |
| Y.YNIDVPSRTARAITTR.S | 1376 | 1391 | 32.94 | 1832.986 | 2.2 | 459.2547 | 4 | 23.02 | 1.10E+08 | 12299 | 2 | |
| Y.YNIDVPSRTAR.A | 1376 | 1386 | 32.15 | 1290.668 | 0.3 | 646.3414 | 2 | 21.08 | 1.24E+08 | 10778 | 5 | |
| Y.NIDVPSRTARAITTR.S | 1377 | 1391 | 27.25 | 1669.922 | 1.6 | 418.4885 | 4 | 20.89 | 2.97E+09 | 10634 | 8 | |
| Y.NIDVPSRTARAI.T | 1377 | 1388 | 25.51 | 1311.726 | -0.4 | 438.249 | 3 | 22.01 | 2.50E+08 | 11510 | 3 | |
| N.IDVPSRTARAITTR.S | 1378 | 1391 | 27.09 | 1555.879 | 0.5 | 389.9773 | 4 | 20.51 | 2.53E+08 | 10338 | 5 | |
| R.SGQTLSK.V | 1392 | 1398 | 25.09 | 719.3813 | -0.5 | 360.6978 | 2 | 12.52 | 2.69E+07 | 4182 | 2 | |

**Supplementary Table S2**

Glycopeptides detected from an 'in-solution' cathepsin G digest of recombinant lubricin (rhPRG4).

Peptides were analyzed with LC-MS/MS, and evaluated manually, as described in Materials and Methods section.

Peptides are listed with their glycan composition, and proposed amino acid attachment sites are underlined (Ser/Thr/Tyr/Asn).

Peptides that occur more than one time in the protein (repeated peptides) are labelled in bold.

Glycopeptides are listed with their adjacent amino acids separated by a dot ('.') . The abbreviation 'X' means that there are more than one possible amino acid option.

| START | END | *m/z* (measured) | Charge | Ret time (min) | Scan no. | PEPTIDE | GLYCAN COMPOSITION HexNAc | Hex | NeuAc | ΔDa | Δppm | CALCULATED NONGLYCOSYLATED PEPTIDE [M+H]+ | Repeated peptides(amino acid positions) |
|---|---|---|---|---|---|---|---|---|---|---|---|---|---|
| 118 | 130 | 742.9028 | 2 | 15.19 | 6233;6370 | K.KAPPPSGASQTIK.S | 1 | 0 | 0 | 0.0028540 | 1.92 | 1281.7161 | |
| 118 | 130 | 495.6037 | 3 | 14.4;14.58 | 5625;5766;5897 | K.KAPPPSGASQTIK.S | 1 | 0 | 0 | 0.0010780 | 0.73 | 1281.7161 | |
| 118 | 130 | 823.9286 | 2 | 13.74 | 5128;5256;5531;6208 | K.KAPPPSGASQTIK.S | 1 | 1 | 0 | 0.0016540 | 1.00 | 1281.7161 | |
| 118 | 130 | 549.6213 | 3 | 14.45 | 5127;5666 | K.KAPPPSGASQTIK.S | 1 | 1 | 0 | 0.0010780 | 0.65 | 1281.7161 | |
| 118 | 130 | 646.6539 | 3 | 15.16 | 6211;6341 | K.KAPPPSGASQTIK.S | 1 | 1 | 1 | 0.0034780 | 1.79 | 1281.7161 | |
| 118 | 130 | 969.4764 | 2 | 15.17 | 6222;6352 | K.KAPPPSGASQTIK.S | 1 | 1 | 1 | 0.0018540 | 0.96 | 1281.7161 | |
| 202 | 219 | 547.1006 | 5 | 13.36 | 4833 | K.DNKKNRTKKKPTPKPPVV.D | 1 | 1 | 1 | 0.0015260 | 0.56 | 2075.2448 | |
| 202 | 220 | 479.4762 | 5 | 12.63 | 4269 | K.DNKKNRTKKKPTPKPPVVD.E | 1 | 0 | 0 | 0.0008260 | 0.35 | 2190.2717 | |
| 202 | 220 | 511.8882 | 5 | 12.52 | 4183 | K.DNKKNRTKKKPTPKPPVVD.E | 1 | 1 | 0 | 0.0080260 | 3.14 | 2190.2717 | |
| 202 | 220 | 712.3813 | 4 | 12.82;12.99;13.17 | 4270;4410;4544;4687 | K.DNKKNRTKKKPTPKPPVVD.E | 1 | 1 | 1 | 0.0041020 | 1.44 | 2190.2717 | |
| 202 | 220 | 570.1049 | 5 | 12.82;13.00;13.17 | 4411;4554;4688 | K.DNKKNRTKKKPTPKPPVVD.E | 1 | 1 | 1 | -0.0038740 | -1.36 | 2190.2717 | |
| 202 | 221 | 671.8676 | 4 | 12.80 | 4398 | K.DNKKNRTKKKPTPKPPVVDE.A | 1 | 1 | 0 | 0.0021020 | 0.78 | 2319.3143 | |
| 202 | 221 | 537.6949 | 5 | 12.65 | 4278 | K.DNKKNRTKKKPTPKPPVVDE.A | 1 | 1 | 0 | -0.0010740 | -0.40 | 2319.3143 | |
| 202 | 221 | 744.6417 | 4 | 13.36 | 4834 | K.DNKKNRTKKKPTPKPPVVDE.A | 1 | 1 | 1 | 0.0031020 | 1.04 | 2319.3143 | |
| 202 | 222 | 762.4017 | 4 | 14.37;14.55 | 5600;5742 | K.DNKKNRTKKKPTPKPPVVDEA.G | 1 | 1 | 1 | 0.0060020 | 1.97 | 2390.3514 | |
| 202 | 223 | 703.8828 | 4 | 13.06 | 4598 | K.DNKKNRTKKKPTPKPPVVDEAG.S | 1 | 1 | 0 | 0.0043020 | 1.53 | 2447.3729 | |
| 202 | 223 | 563.3073 | 5 | 13.02;13.20 | 4570;4713 | K.DNKKNRTKKKPTPKPPVVDEAG.S | 1 | 1 | 0 | 0.0023260 | 0.83 | 2447.3729 | |
| 202 | 223 | 1035.2064 | 3 | 14.29 | 5543 | K.DNKKNRTKKKPTPKPPVVDEAG.S | 1 | 1 | 1 | 0.0041780 | 1.35 | 2447.3729 | |
| 202 | 223 | 776.6573 | 4 | 14.45 | 5665;5938;6075 | K.DNKKNRTKKKPTPKPPVVDEAG.S | 1 | 1 | 1 | 0.0069020 | 2.22 | 2447.3729 | |
| 202 | 226 | 1120.9166 | 3 | 17.62 | 8102 | K.DNKKNRTKKKPTPKPPVVDEAGSGL.D | 1 | 1 | 1 | -0.0027220 | -0.81 | 2704.5104 | |
| 205 | 220 | 623.0892 | 4 | 12.86 | 4448 | K.KNRTKKKPTPKPPVVD.E | 1 | 1 | 1 | 0.0005020 | 0.20 | 1833.1069 | |
| 205 | 220 | 498.6725 | 5 | 12.90 | 4477 | K.KNRTKKKPTPKPPVVD.E | 1 | 1 | 1 | -0.0010740 | -0.43 | 1833.1069 | |
| 205 | 223 | 916.1514 | 3 | 14.38 | 5612 | K.KNRTKKKPTPKPPVVDEAG.S | 1 | 1 | 1 | 0.0040780 | 1.48 | 2090.2080 | |
| 205 | 223 | 687.3655 | 4 | 14.31;14.84 | 5559;5964 | K.KNRTKKKPTPKPPVVDEAG.S | 1 | 1 | 1 | 0.0046020 | 1.68 | 2090.2080 | |
| 206 | 220 | 623.3268 | 4 | 14.61 | 5788 | K.NRTKKKPTPKPPVVDE.A | 1 | 1 | 1 | 0.0033020 | 1.33 | 1834.0545 | |
| 206 | 223 | 873.4525 | 3 | 14.91 | 6016 | K.NRTKKKPTPKPPVVDEAG.S | 1 | 1 | 1 | 0.0022780 | 0.87 | 1962.1131 | |
| 206 | 223 | 655.3436 | 4 | 14.98 | 6072 | K.NRTKKKPTPKPPVVDEAG.S | 1 | 1 | 1 | 0.0119020 | 4.55 | 1962.1131 | |
| 206 | 223 | 524.4744 | 5 | 14.98 | 6074 | K.NRTKKKPTPKPPVVDEAG.S | 1 | 1 | 1 | 0.0022260 | 0.85 | 1962.1131 | |
| 208 | 223 | 835.4384 | 3 | 14.98 | 6076 | N.RTKKKPTPKPPVVDEAG.S | 1 | 1 | 1 | 0.0028780 | 1.15 | 1848.0702 | |
| 208 | 223 | 626.8303 | 4 | 14.98 | 6073 | N.RTKKKPTPKPPVVDEAG.S | 1 | 1 | 1 | 0.0016020 | 0.64 | 1848.0702 | |
| 210 | 220 | 470.2731 | 3 | 15.11 | 6179 | K.KKPTPKPPVVD.E | 1 | 0 | 0 | 0.0001780 | 0.13 | 1205.7252 | |
| 210 | 220 | 621.3227 | 3 | 15.97 | 6821 | K.KKPTPKPPVVD.E | 1 | 1 | 1 | 0.0007780 | 0.42 | 1205.7252 | |
| 210 | 221 | 664.3367 | 3 | 16.48 | 7217 | K.KKPTPKPPVVDE.A | 1 | 1 | 1 | 0.0001780 | 0.09 | 1334.7678 | |
| 210 | 223 | 707.0245 | 3 | 16.80 | 7464 | K.KKPTPKPPVVDEAG.S | 1 | 1 | 1 | 0.0049780 | 2.35 | 1462.8264 | |
| 252 | 267 | 791.4083 | 3 | 21.59 | 11178 | K.ITTAKPINPRPSLPPN.S | 1 | 1 | 1 | 0.0024780 | 1.04 | 1715.9803 | |
| 252 | 267 | 913.1203 | 3 | 21.26 | 10921 | K.ITTAKPINPRPSLPPN.S | 2 | 2 | 1 | 0.0063080 | 2.30 | 1715.9803 | |
| 252 | 272 | 964.1537 | 3 | 20.44 | 10287 | K.ITTAKPINPRPSLPPNSDTSK.E | 1 | 1 | 1 | 0.0050780 | 1.76 | 2234.2139 | |
| 252 | 272 | 723.3666 | 4 | 20.25 | 10141 | K.ITTAKPINPRPSLPPNSDTSK.E | 1 | 1 | 1 | 0.0031020 | 1.07 | 2234.2139 | |
| 252 | 272 | 814.6498 | 4 | 20.30 | 10181 | K.ITTAKPINPRPSLPPNSDTSK.E | 2 | 2 | 1 | 0.0037320 | 1.15 | 2234.2139 | |
| 316 | 326 | 495.2504 | 3 | 15.72;16.24 | 6638;6499 | K.TSAKDLAPTSK.V | 1 | 1 | 0 | -0.0007220 | -0.49 | 1118.6052 | |
| 316 | 326 | 592.2832 | 3 | 16.24 | 7831 | K.TSAKDLAPTSK.V | 1 | 1 | 1 | 0.0022780 | 1.28 | 1118.6052 | |
| 316 | 327 | 625.3057 | 3 | 18.60 | 8864 | K.TSAKDLAPTSKV.L | 1 | 1 | 1 | 0.0013780 | 0.74 | 1217.6736 | |
| 316 | 334 | 703.8680 | 4 | 18.53 | 8813 | K.TSAKDLAPTSKVLAKPTPK.A | 2 | 1 | 1 | 0.0053320 | 1.90 | 1953.1379 | |
| 316 | 334 | 817.1553 | 4 | 18.97 | 9145 | K.TSAKDLAPTSKVLAKPTPK.A | 2 | 2 | 2 | 0.0063320 | 1.94 | 1953.1379 | |
| 317 | 326 | 494.5906 | 3 | 16.77 | 7441 | T.SAKDLAPTSKV.L | 1 | 1 | 0 | -0.0008220 | -0.55 | 1116.63 | |
| 317 | 326 | 591.6233 | 3 | 17.64 | 8120 | T.SAKDLAPTSKV.L | 1 | 1 | 1 | 0.0018780 | 1.06 | 1116.63 | |
| 317 | 334 | 719.1201 | 4 | 17.60 | 8090 | T.SAKDLAPTSKVLAKPTPK.A | 2 | 2 | 1 | 0.0086320 | 3.00 | 1852.09 | |
| 320 | 326 | 694.3148 | 2 | 15.55 | 6508;6562 | K.DLAPTSK.V | 1 | 1 | 1 | 0.0013540 | 0.98 | 731.39 | |
| 320 | 334 | 493.7768 | 4 | 18.51 | 8788 | K.DLAPTSKVLAKPTPK.A | 2 | 0 | 0 | 0.0005320 | 0.27 | 1565.9261 | |
| 320 | 334 | 863.1029 | 3 | 18.67 | 8916 | K.DLAPTSKVLAKPTPK.A | 2 | 2 | 1 | 0.0083080 | 3.21 | 1565.9261 | |
| 320 | 334 | 647.5783 | 4 | 18.71 | 8943 | K.DLAPTSKVLAKPTPK.A | 2 | 2 | 1 | 0.0055320 | 2.14 | 1565.9261 | |
| 320 | 334 | 607.0639 | 4 | 19.20 | 9321 | K.DLAPTSKVLAKPTPK.A | 2 | 1 | 1 | 0.0007320 | 0.30 | 1565.9261 | |
| 326 | 333 | 406.8990 | 3 | 13.70 | 5099;6852 | K.VLAKPTPK.A | 1 | 1 | 0 | -0.0003220 | -0.26 | 853.5506 | |
| 326 | 333 | 503.9305 | 3 | 16.00 | 6848 | K.VLAKPTPK.A | 1 | 1 | 1 | -0.0012220 | -0.81 | 853.5506 | |
| 326 | 333 | 755.3931 | 2 | 16.01 | 6852 | K.VLAKPTPK.A | 1 | 1 | 1 | 0.0007540 | 0.50 | 853.5506 | |
| 327 | 333 | 470.9079 | 3 | 15.31 | 6322 | V.LAKPTPK.A | 1 | 1 | 1 | -0.0006220 | -0.44 | 754.4822 | |
| 402 | 410 | 542.2612 | 3 | 15.11 | 6174 | **X.KEPAPTTPK.X** | 1 | 1 | 1 | 0.0003780 | 0.23 | 968.5411 | 402-10;433-41;449-57;472-80;496-504;558-66;566-74;582-90;606-14;678-86;686-94;694-702;718-26;762-71;771-79;787-95;832-40 |
| 402 | 410 | 812.8888 | 2 | 15.11 | 6178 | **X.KEPAPTTPK.X** | 1 | 1 | 1 | 0.0016540 | 1.02 | 968.5411 | |
| 615 | 622 | 524.2638 | 2 | 15.71 | 6627 | **K.ETAPTTPK.X** | 1 | 0 | 0 | -0.0001460 | -0.14 | 844.4411 | 615-622;703-710;825-832 |
| 615 | 622 | 605.2905 | 2 | 15.19 | 6236 | **K.ETAPTTPK.X** | 1 | 1 | 0 | 0.0004540 | 0.38 | 844.4411 | 615-622;703-710;825-832 |
| 615 | 622 | 750.8367 | 2 | 16.43 | 7174 | **K.ETAPTTPK.X** | 1 | 1 | 1 | -0.0025460 | -1.70 | 844.4411 | 615-622;703-710;825-832 |
| 718 | 731 | 423.9937 | 4 | 13.37;13.54;13.71;14.07 | 4974;5373;5109;4839 | **L.KEPAPTTPKKPAPK.E** | 1 | 0 | 0 | -0.0000980 | -0.06 | 1489.8737 | 718-731;787-800 |
| 718 | 731 | 564.9890 | 3 | 13.75;14.56;14.74 | 5140;5755;5888 | **L.KEPAPTTPKKPAPK.E** | 1 | 0 | 0 | -0.0006220 | -0.37 | 1489.8737 | 718-731;787-800 |
| 718 | 731 | 464.5075 | 4 | 12.99;13.17;14.64 | 4545;4686;5843 | **L.KEPAPTTPKKPAPK.E** | 1 | 1 | 0 | 0.0023020 | 1.24 | 1489.8737 | 718-731;787-800 |
| 718 | 731 | 619.0070 | 3 | 13.02;13.20 | 4573;4714 | **L.KEPAPTTPKKPAPK.E** | 1 | 1 | 0 | -0.0025460 | 0.31 | 1489.8737 | 718-731;787-800 |
| 718 | 731 | 1073.5551 | 2 | 14.68 | 5840 | **L.KEPAPTTPKKPAPK.E** | 1 | 1 | 1 | 0.0016540 | 0.77 | 1489.8737 | 718-731;787-800 |
| 718 | 731 | 716.0397 | 3 | 14.62;14.79 | 5799;5925 | **L.KEPAPTTPKKPAPK.E** | 1 | 1 | 1 | 0.0032780 | 1.53 | 1489.8737 | 718-731;787-800 |
| 718 | 731 | 537.2816 | 4 | 14.41;14.59;14.78 | 5638;5778;5912 | **L.KEPAPTTPKKPAPK.E** | 1 | 1 | 1 | 0.0033020 | 1.54 | 1489.8737 | 718-731;787-800 |
| 718 | 731 | 430.0257 | 5 | 14.59;14.78 | 5779;5913 | **L.KEPAPTTPKKPAPK.E** | 1 | 1 | 1 | -0.0018740 | -0.87 | 1489.8737 | 718-731;787-800 |
| 718 | 731 | 474.7636 | 4 | 13.44;13.80;13.88 | 4898;5177;5229 | **L.KEPAPTTPKKPAPK.E** | 2 | 0 | 0 | 0.0001320 | 0.07 | 1489.8737 | 718-731;787-800 |
| 718 | 731 | 515.2770 | 4 | 13.62;13.97 | 5039;5297 | **L.KEPAPTTPKKPAPK.E** | 2 | 1 | 0 | 0.0009320 | 0.45 | 1489.8737 | 718-731;787-800 |
| 718 | 731 | 588.0514 | 4 | 14.83 | 5952 | **L.KEPAPTTPKKPAPK.E** | 2 | 1 | 1 | 0.0031320 | 1.33 | 1489.8737 | 718-731;787-800 |
| 718 | 731 | 628.5643 | 4 | 14.26;14.62 | 5523;5801 | **L.KEPAPTTPKKPAPK.E** | 2 | 2 | 1 | 0.0019320 | 0.77 | 1489.8737 | 718-731;787-800 |
| 718 | 731 | 837.7499 | 3 | 14.36 | 5799 | **L.KEPAPTTPKKPAPK.E** | 2 | 2 | 1 | 0.0017080 | 0.68 | 1489.8737 | 718-731;787-800 |
| 718 | 731 | 934.7829 | 3 | 14.78;14.96 | 5915;6055 | **L.KEPAPTTPKKPAPK.E** | 2 | 2 | 2 | 0.0053080 | 1.89 | 1489.8737 | 718-731;787-800 |
| 718 | 731 | 701.3372 | 4 | 14.71;14.87;15.04;15.22 | 5851;5990;6124;6262 | **L.KEPAPTTPKKPAPK.E** | 2 | 2 | 2 | -0.0018680 | -0.67 | 1489.8737 | 718-731;787-800 |
| 718 | 731 | 561.2717 | 5 | 14.89 | 6004 | **L.KEPAPTTPKKPAPK.E** | 2 | 2 | 2 | 0.0005560 | 0.20 | 1489.8737 | 718-731;787-800 |
| 871 | 887 | 1089.5138 | 2 | 19.57 | 9611 | H.KSPDESTPELSAEPTPK.A | 1 | 1 | 0 | 0.0019540 | 0.90 | 1812.8862 | |
| 871 | 887 | 1235.0596 | 2 | 20.34 | 10209 | H.KSPDESTPELSAEPTPK.A | 1 | 1 | 1 | -0.0018460 | -0.75 | 1812.8862 | |
| 871 | 887 | 823.7111 | 3 | 20.24 | 10128 | H.KSPDESTPELSAEPTPK.A | 1 | 1 | 1 | 0.0049780 | 2.02 | 1812.8862 | |
| 880 | 887 | 749.8493 | 2 | 17.19 | 7763 | E.LSAEPTPK.A | 1 | 1 | 1 | 0.0019540 | 1.30 | 842.4618 | |
| 882 | 887 | 649.7911 | 2 | 13.50 | 4947 | S.AEPTPK.A | 1 | 1 | 1 | 0.0016540 | 1.27 | 642.3457 | |
| 1073 | 1087 | 932.1101 | 3 | 16.19 | 6992 | M.LQTTTRPNQTPNSKL.V | 3 | 3 | 0 | 0.0059380 | 2.13 | 1698.9133 | |
| 1073 | 1087 | 1029.1414 | 3 | 17.12 | 7711 | M.LQTTTRPNQTPNSKL.V | 3 | 3 | 1 | 0.0044380 | 1.44 | 1698.9133 | |
| 1073 | 1087 | 1126.1714 | 3 | 17.72;17.90;18.07 | 8179;8315;8449 | M.LQTTTRPNQTPNSKL.V | 3 | 3 | 2 | -0.0009620 | -0.28 | 1698.9133 | |
| 1073 | 1087 | 1223.2062 | 3 | 18.54 | 8812 | M.LQTTTRPNQTPNSKL.V | 3 | 3 | 3 | 0.0080380 | 2.19 | 1698.9133 | |
| 1073 | 1087 | 917.6573 | 4 | 18.57 | 8838 | M.LQTTTRPNQTPNSKL.V | 3 | 3 | 3 | 0.0113620 | 3.10 | 1698.9133 | |
| 1073 | 1087 | 837.7497 | 3 | 16.77 | 7438 | M.LQTTTRPNQTPNSKL.V | 4 | 0 | 0 | 0.0037680 | 1.50 | 1698.9133 | |
| 1073 | 1087 | 891.7679 | 3 | 16.69 | 7373 | M.LQTTTRPNQTPNSKL.V | 4 | 1 | 0 | 0.0055680 | 2.08 | 1698.9133 | |
| 1073 | 1087 | 1042.8168 | 3 | 17.42 | 7945 | M.LQTTTRPNQTPNSKL.V | 4 | 2 | 1 | 0.0040680 | 1.30 | 1698.9133 | |
| 1073 | 1087 | 1139.8500 | 3 | 18.32 | 8643 | M.LQTTTRPNQTPNSKL.V | 4 | 2 | 2 | 0.0082680 | 2.42 | 1698.9133 | |
| 1073 | 1087 | 1096.8350 | 3 | 16.90 | 7542 | M.LQTTTRPNQTPNSKL.V | 4 | 3 | 1 | 0.0058680 | 1.78 | 1698.9133 | |
| 1073 | 1087 | 895.6526 | 4 | 17.74 | 8192 | M.LQTTTRPNQTPNSKL.V | 4 | 3 | 2 | 0.0085920 | 2.40 | 1698.9133 | |
| 1073 | 1087 | 1290.8986 | 3 | 18.40 | 8708 | M.LQTTTRPNQTPNSKL.V | 4 | 3 | 3 | 0.0058680 | 1.52 | 1698.9133 | |
| 1073 | 1087 | 968.4262 | 4 | 18.39 | 8696 | M.LQTTTRPNQTPNSKL.V | 4 | 3 | 3 | 0.0075920 | 1.96 | 1698.9133 | |
| 1122 | 1130 | 880.8947 | 2 | 25.61 | 14311 | D.MDYLPRVPN.Q | 1 | 1 | 1 | 0.0039540 | 2.25 | 1104.5506 | |
| 1122 | 1130 | 684.6301 | 3 | 25.69 | 14372 | D.MDYLPRVPN.Q | 1 | 1 | 2 | 0.0021780 | 1.06 | 1104.5506 | |
| 1122 | 1130 | 1026.4418 | 2 | 25.72 | 14397 | D.MDYLPRVPN.Q | 1 | 1 | 2 | 0.0027540 | 1.34 | 1104.5506 | |
| 1158 | 1165 | 1047.4613 | 2 | 24.38 | 13357 | L.RNGTVLAF.R | 2 | 5 | 0 | 0.0025840 | 1.23 | 877.4900 | |
| 1158 | 1165 | 1128.4889 | 2 | 24.20 | 13213 | L.RNGTVLAF.R | 2 | 6 | 0 | 0.0049840 | 2.21 | 877.4900 | |
